# Metastable Neural Assemblies on a Wiring–Weight Continuum

**DOI:** 10.64898/2026.03.16.712138

**Authors:** Felix J. Schmitt, David Dahmen, Franziska L. Müller, Martin P. Nawrot

## Abstract

Neural population activity typically evolves on low-dimensional manifolds and can be described as trajectories through quasi-stable assembly states. Here we develop a unified definition of clustered neural networks with local excitatory–inhibitory balance in which enhanced within-cluster effective coupling is implemented through connection probability (structural clustering), synaptic efficacy (weight clustering), or any mixture of both. We introduce a single mixing parameter *κ* ∈ [0, 1] that redistributes a defined cluster contrast between connection probabilities and synaptic efficacies while preserving the mean input of the underlying balanced random network. Mean-field theory and binary-network simulations show that metastable dynamics are supported across the full *κ* continuum. Varying *κ* changes higher-order input structure, reshaping multistable regimes, correlation structure, and the balance between single- and multi-cluster episodes. Because real nervous systems jointly organize topology and synaptic strength, our approach provides a biologically interpretable parametrization of clustered assembly models and a basis for future models combining structural and functional plasticity. We further demonstrate metastable switching in spiking leaky integrate-and-fire networks and in a BrainScaleS-2 neuromorphic implementation for the cases of structural, weight, and combined clustering. The *κ*-framework offers a controlled translation axis for neuromorphic and other constrained substrates, exposing trade-offs between routed synapse count, fan-in, synaptic weight resolution, and calibration when implementing attractor-based computational primitives.

## Introduction

Across species, neuronal population activity often exhibits recurring dynamical motifs in low-dimensional state space [1–7], for example continuous-attractor dynamics, categorical fixed-point dynamics that yield winner-take-all (WTA) outcomes, and sequences of quasi-stable assembly states (metastability). These dynamical motifs in state space should be distinguished from structural motifs in attractor network models, which specify how recurrent coupling is instantiated in a circuit. Continuous attractor neural network (CANN) models, for example, realize a low-dimensional attractor manifold through approximately translation-invariant (for example ring-structured) recurrent coupling. In rodents, grid-cell representations have been modeled as 2D continuous attractors [8], and the head-direction system is a canonical case for ring-attractor dynamics [9]. Analogously, in insects the anatomy of the central complex provides an explicitly ring-like implementation of a CANN that encodes heading direction and supports angular path integration during locomotion [10, 11]. Categorical fixed-point (WTA) dynamics can be implemented by recurrent competition between choice-selective excitatory pools mediated through shared or lateral feedback inhibition, as in biophysically grounded decision-circuit models [12–17].

In the mammalian neocortex, ongoing activity is often non-stationary and, during the performance of a task, unfolds as sequences of quasi-stable neuronal population states [18–23]. Natural candidate substrates for such metastability are neuronal assemblies, subsets of neurons that tend to co-activate and can support pattern completion, persistent activity, and switching between activity states [7, 24, 25]. Structurally, assemblies are thought to be promoted by non-random recurrent connectivity in cortical microcircuits, including clustered and fine-scale subnetwork organization [26–30]. Here we use assembly to refer to the dynamical co-activation pattern [7], and clustering to refer to the underlying non-random connectivity that can bias such patterns.

Consistent with this picture, excitation–inhibition balanced network models [31, 32] with embedded clusters of recurrently coupled excitatory neurons exhibit slow switching among different assembly patterns and reproduce key signatures of metastability [33–36]. In such models, trial-to-trial variability is not merely noise but reflects transitions between latent states and induces state-dependent modulation of neuronal correlations [23, 33, 37, 38]. Computation can therefore be viewed as trajectories through a low-dimensional state space [39,40], supporting robust coding with flexible reconfiguration and enabling cognitive functions such as action selection, decision-making, and working memory [23, 41, 42].

Clustered network topology can be realized by distinct architectures, including increased within-cluster connection probability (structural clustering) [35], increased within-cluster synaptic efficacy (weight clustering) [33, 35], or combinations thereof [34]. Extensions incorporate the locally balanced excitatory-inhibitory (EI) cluster organization [23, 36] and the hierarchical cluster structure [35], and plasticity models can drive the emergence of clustered organization [43, 44]. Related work likewise shows that structured connectivity can shape network operating regimes and computation [45]. Importantly, this architectural diversity is not merely a modeling choice but reflects substrate constraints. Biological circuits face constraints on wiring, sparsity, and synapse statistics [46, 47], whereas artificial substrates, especially neuromorphic platforms, often impose explicit restrictions on fan-in and fan-out, routing, and weight resolution [48–51]. Consequently, different substrates naturally privilege different ways of realizing effective within-versus across-cluster coupling.

To enable a controlled comparison of alternative implementations of clustered recurrent coupling, we introduce a unified network family parameterized by a single mixing parameter *κ* ∈ [0, 1]. The parameter continuously redistributes cluster contrast between connection probabilities (*κ* = 0, structural clustering) and synaptic efficacies (*κ* = 1, weight clustering), with intermediate values representing mixed implementations. In contrast to previous studies that treated these architectures separately or in selected combinations [33–35], our formulation places them on a common continuum while preserving the effective within-versus across-cluster coupling contrast and the expected mean recurrent input of the underlying balanced random network. This allows us to isolate the dynamical consequences of how a matched mesoscale coupling structure is implemented. Although first-order recurrent input is preserved, varying *κ* changes expected indegree, input variance, and higher-order input statistics, and thereby reshapes fixed-point structure, neuronal correlations, and the balance between single- and multi-cluster episodes. We characterize these effects using mean-field theory and microscopic binary-network simulations, and then test their qualitative robustness in spiking leaky integrate-and-fire networks and on spiking neuromorphic hardware. Together, these results show that metastable dynamics are supported across the full wiring–weight continuum in binary, spiking, and neuromorphic implementations, although the corresponding operating regimes and switching patterns depend on *κ*.

### Unified network model

We define a family of networks with embedded excitatory-inhibitory (EI) clusters by modifying the standard balanced random network as schematically depicted in Figure 1a. Clustering of neurons is achieved by redistribution of synaptic coupling. We define a single mixing parameter *κ* that determines the ratio of structural clustering versus weight clustering (Figure 1b). The redistribution of synaptic coupling is designed to maintain the balanced mean input condition of the balanced random network. Figure 1c illustrates the desired dynamic regime of metastability for the endpoints of either purely structural (c1) or purely weight clustering (c2). Here, we present the model construction, whereas details of the simulations and mean-field analysis are provided in the Methods.

**Figure 1.**
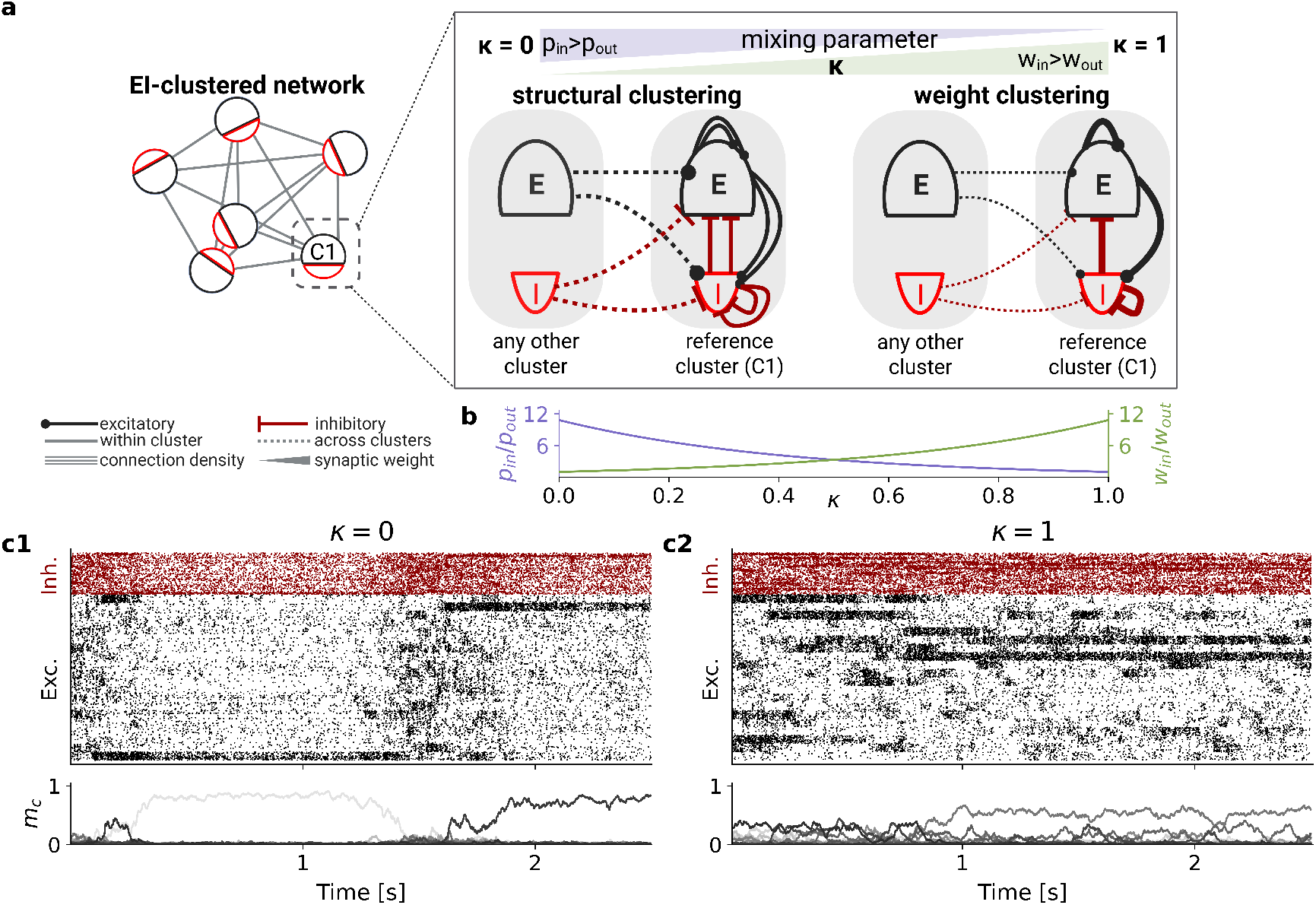
Unified EI clustering with mixing parameter κ produces metastable switching at both endpoints of pure structural clustering versus pure weight clustering. (a) Schematic of an EI-clustered network with NQ clusters. The mixing parameter κ determines the ratio of structural clustering and weight clustering for excitatory and inhibitory connections while keeping the cluster contrast fixed. (b) The example shown indicates how κ scales connection probabilities within (p_in_) and across clusters (p_out_) versus weight parameters within (w_in_) and across clusters (w_out_) for excitatory projections, using network parameters N_Q_ = 20 and R_E+_ = 7.25. (c) Binary network simulations (N_Q_ = 20, R_E+_ = 7.25, R_j_ = 0.8) are shown as an event raster (top) and mean excitatory per-cluster activities m_c_ = m_E,c_ (bottom) for structural clustering (κ = 0, panel c1) and weight clustering (κ = 1, panel c2). Both regimes exhibit metastable dynamics. Pure structural clustering (κ = 0) shows larger single-cluster activations and fewer multi-cluster episodes. Events indicate 0→1 state transitions in the binary units; 2.5 s corresponds to ∼ 250 asynchronous update cycles (see Methods).

### Balanced random network

We consider a network of *N* neurons (*N*_E_ excitatory, *N*_I_ inhibitory). A neuron of type *β* ∈ {E, I} connects to a neuron of type *α* ∈ { E, I } with probability *p*_*αβ*_. To maintain the balanced-state regime, synaptic efficacies are scaled with network size as

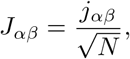

where *j*_*α β*_ is constant, independent of *N* [36]. The 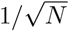 scaling ensures that the mean excitatory and inhibitory inputs can cancel while the variance of the total synaptic input remains constant with network size [31]. Size-independent efficacies are set by

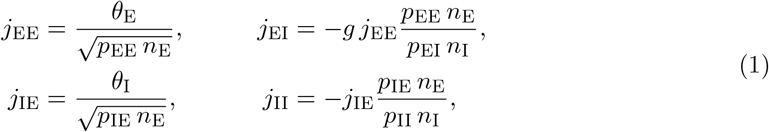

where *n*_*β*_ = *N*_*β*_*/N, θ*_*α*_ are thresholds and *g* sets inhibitory strength.

### Mixed structural- and weight clustering

Neurons are partitioned into *N*_*Q*_ non-overlapping clusters, each containing *N*_*β*_*/N*_*Q*_ neurons of type *β* ∈ E, I . We introduce a mixing parameter *κ* ∈ [0, 1] that redistributes cluster contrast between connection probabilities and synaptic weights. For a neuron pair (*i, j*), we define the projection-specific contrast *R*_*ij*_ based on the types of the pre- and postsynaptic neurons (*α*_*j*_, *α*_*i*_):

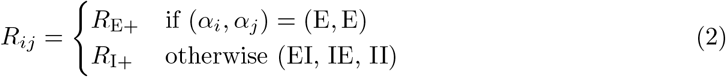

The clustered connection probability *p*_*ij*_ and synaptic weight *J*_*ij*_ are then given by

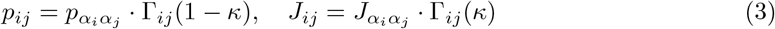

where the contrast function Γ_*ij*_(*x*) modulates the baseline according to cluster indices *m*_*i*_, *m*_*j*_:

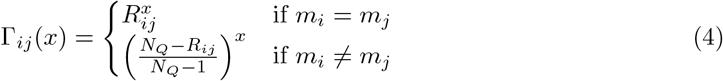

This formulation preserves the expected mean recurrent input of the underlying balanced random network for all *κ*, because the product of connection-probability and synaptic-weight modulation keeps the effective within-versus across-cluster coupling contrast fixed. The expected indegree, however, is not generally preserved for intermediate *κ*. For a given cluster contrast *R* ∈ {*R*_E+_, *R*_I+_} and number of clusters *N*_*Q*_, the projection-specific expected indegree relative to the balanced random network is

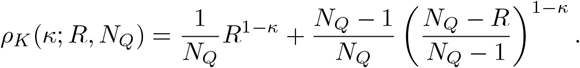

Thus, *ρ*_*K*_ = 1 at the endpoints *κ* = 0 and *κ* = 1, but can be substantially smaller at intermediate *κ*, particularly for large cluster contrasts. Varying *κ* therefore changes not only whether cluster contrast is implemented through connection probabilities or synaptic weights, but also the expected number of realized contacts and higher input moments. Accordingly, the parameter *κ* should be interpreted as an effective implementation coordinate: small *κ* values express clustered coupling mainly through connection or contact statistics, large *κ* values mainly through synaptic efficacy, and intermediate values implement mixtures of both while maintaining the same effective within-cluster coupling. Thus, *κ* does not specify a unique microscopic biological process, but defines a controlled axis for comparing circuits or substrates that realize similar mesoscale coupling through different combinations of wiring and synaptic strength. Inhibitory clustering is linked to excitatory clustering through *R*_I+_ = 1 + *R*_*j*_(*R*_E+_ − 1), where *R*_*j*_ ∈ [0, 1] controls the relative degree of inhibitory organization.

### Realization of the clustered multigraph

To account for the likely case where within-cluster probabilities might exceed unity, we realize the connectivity as a directed Poisson multigraph and not as an Erdős–Rényi graph with Bernoulli edges without multiple contacts that is commonly used [52]. Synapse multiplicities *K*_*ij*_ for all ordered pairs (including *i* = *j*, autapses) are drawn independently:

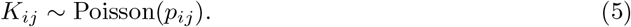

For *β* ∈ {E, I} and *s* ∈ {1, …, *N*_*Q*_}, define

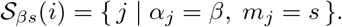

Then

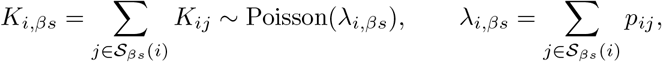

so that

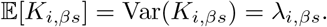

The eigenvalue spectrum and an alternative fixed indegree construction are reported in the Supplementary Information.

## Results

### Mean-field analysis predicts metastable regimes

To map the parameter space where the clustered network family supports multiple coexisting macroscopic states as a function of *κ*, we analyze a simplified mean-field description of the clustered binary network (Fig. 2). Using the effective response function (ERF) approach [53], we locate fixed points of the reduced macroscopic dynamics under a Gaussian approximation for recurrent input, neglecting temporal correlations (Methods). Across the three representative positions along the *κ* continuum, increasing the excitatory clustering strength *R*_E+_ can induce bifurcations that create regions with multiple admissible fixed points for the same *R*_E+_ (Fig. 2a–e). However, for *κ* = 1 and at a lower average connection probability 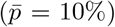, the mean-field approximation predicts fixed points throughout this regime without a bifurcation point and corresponding multistable structure (Fig. 2f). The coexistence of well-separated stable solutions is consistent with a metastable regime in finite networks [39], where intrinsic fluctuations can drive switching between population states. The location and geometry of these multistable regions shift markedly with *κ*. Matching *R*_E+_ across different mixtures thus does not, in general, match the macroscopic operating point.

**Figure 2.**
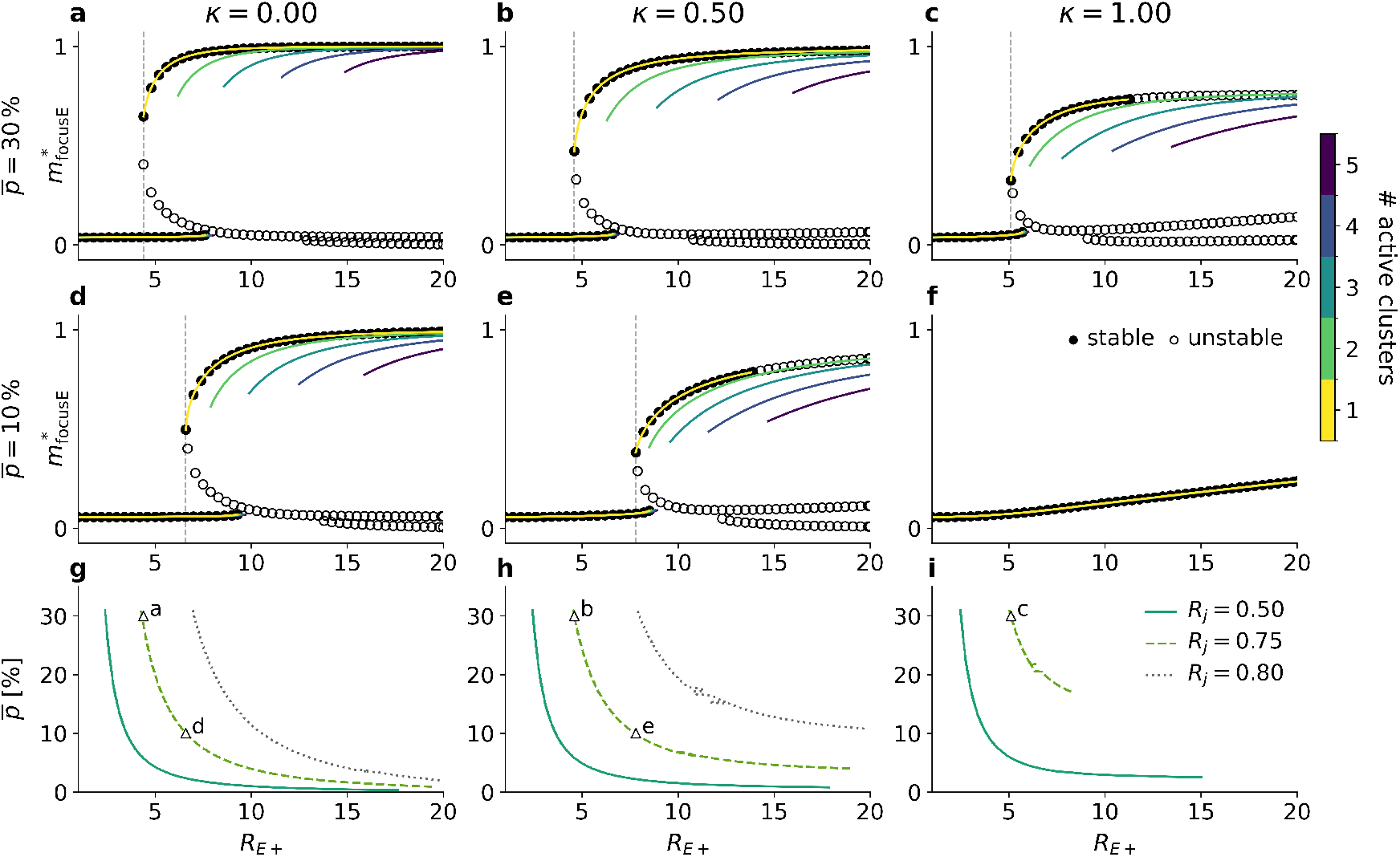
Mean-field fixed-point structure and onset of multistability depend on mixing parameter κ. (a–c) Predicted fixed points of the focused excitatory cluster activity 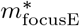 as a function of excitatory clustering strength RE_+_ at baseline connection probability across all neurons of 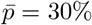, shown for (a) κ = 0, (b) κ = 0.5, and (c) κ = 1. R_j_ = 0.75 is fixed in all cases. Colored branches correspond to solutions with different numbers of simultaneously active clusters (stable branches shown). Black markers indicate stable (filled) and unstable (open) single-active-cluster fixed points. (d–f) Same as (a–c) for 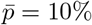 Vertical markers indicate the first appearance of clearly separated coexisting solutions for a given RE+. (g–i) Corresponding first bifurcation boundary in the 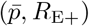 plane; metastability is possible to the right of the boundary, while only the global mode exists to the left. Branches are discontinued if no bifurcation was found (f). All analyses were performed using the quenched closure and a root-finding solver.

A key difference is a trade-off between activity levels and the required baseline connection probability 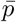. For structural clustering (*κ* = 0), multistability appears already at lower 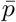, but the corresponding activity 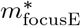of active clusters increases strongly and can reach saturation. For weight clustering (*κ* = 1), multiple stable branches emerge at lower 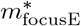 for comparable *R*_E+_, but may require a higher 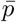 for the first bifurcation. Intermediate mixtures (e.g. *κ* = 0.5) interpolate between these extremes, yet still show distinct shifts in the fixed-point structure. Thus, redistributing contrast between *p* and *J* preserves the expected mean recurrent input, but changes expected indegree, input variance, and higher-order input statistics that affect the bifurcation structure in the mean-field description.

To summarize these effects in a compact regime diagram, we extract the first bifurcation boundary in the 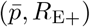 plane (Fig. 2g–i). The region to the right of the boundary admits multiple stable solutions, hence metastability is possible, whereas to the left only the homogeneous global mode exists. Because the analysis omits noise-driven transitions, finite-size effects, and the basin geometry around each fixed point, it cannot predict switching rates or guarantee that coexisting fixed points are dynamically visited in simulations. We therefore interpret the boundary as a qualitative map of how *κ* reshapes the multistable regime, rather than a quantitatively exact phase boundary. The corresponding fixed indegree analysis is shown in Supplementary Fig. S2.

### Binary simulations reproduce metastability across *κ*

We next test whether the *κ*-dependent differences predicted by mean-field theory carry over to microscopic dynamics by simulating the binary network at fixed excitatory clustering strength *R*_E+_ while varying the clustering mixture *κ* (Fig. 3). All three mixtures exhibit metastable switching, but the dominant switching pattern depends on *κ*. For structural clustering (*κ* = 0), trajectories are dominated by long-lived single-cluster up-states interrupted by brief excursions. For intermediate mixing (*κ* = 0.5), switching between single-cluster states becomes more frequent. For weight clustering (*κ* = 1), trajectories more frequently visit states with multiple simultaneously active clusters as previously reported [36].

**Figure 3.**
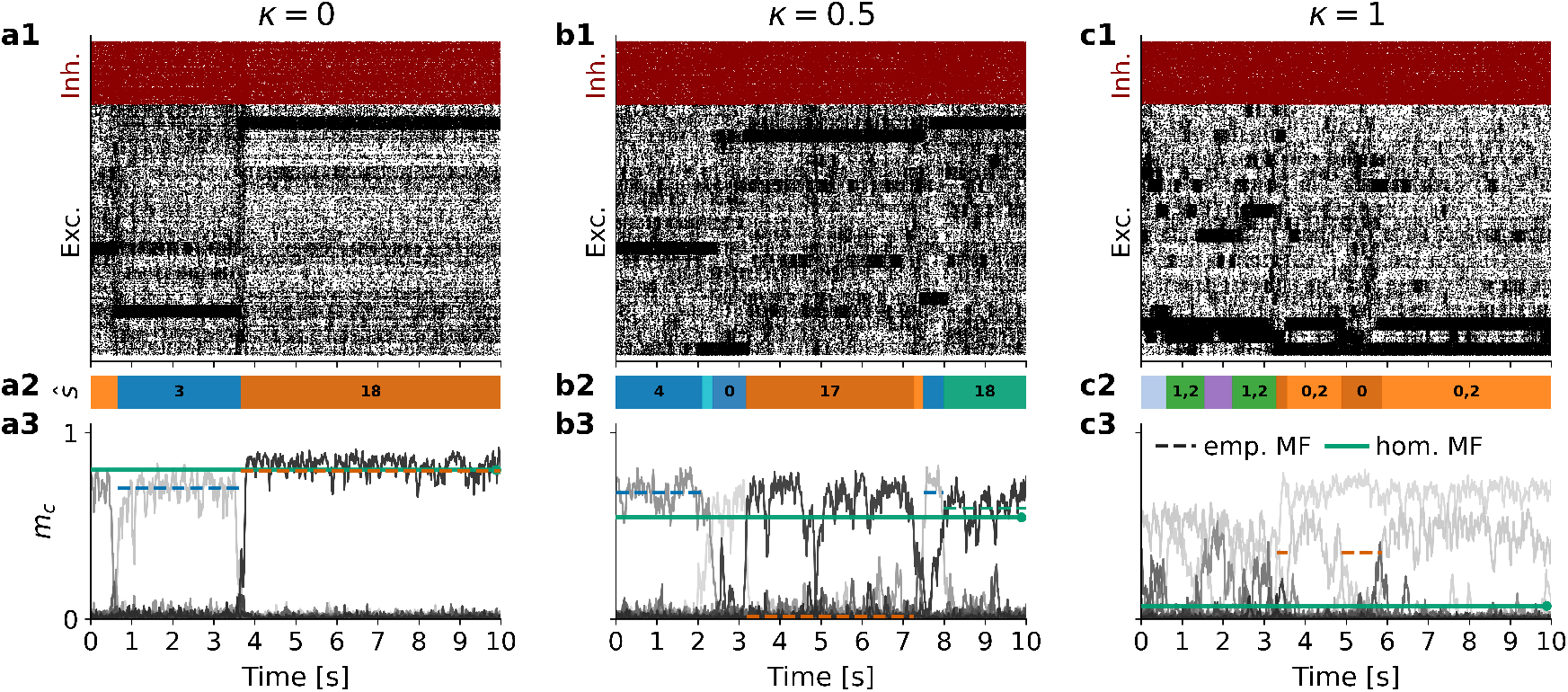
Binary network metastable regimes differ across clustering mixtures. Binary networks (N_E_ = 8,000, N_I_ = 2,000, N_Q_ = 20) were simulated at fixed R_E+_ = 7.25 and R_j_ = 0.79 for (a) κ = 0, (b) κ = 0.5, and (c) κ = 1. Top (a1–c1): rasters of onset events illustrate metastable switching with κ-dependent regimes (predominantly single-cluster states with long dwell times for κ = 0; more frequent switching for κ = 0.5; more multi-cluster episodes for κ = 1). Middle (a2–c2): inferred states ŝ, represented as sets of up to two active clusters. Bottom (a3–c3): mean cluster activity mc. Solid horizontal lines show standard mean-field predictions based on theoretical connectivity expectations; dashed horizontal lines show empirical mean-field predictions based on cluster- and projection-specific connectivity moments measured from the corresponding realization. Both predict single-active-cluster fixed points. Near the multistability boundary, mean-field predictions and simulations can diverge, notably in b3 and c3. Predictions were computed using the quenched closure and damped Picard iteration.

Fig. 3 specifically shows scenarios that are close to the boundary of the multistable regime. At this point, fluctuations are particularly strong, driving the system to transition between various high-activity states. Farther inside the multistable regime, as obtained by reducing *R*_*j*_ from 0.79 to 0.75 (Fig. 4) or by increasing *R*_E+_, (Fig. 2 and Supplementary Table 1) switching is substantially reduced. Because the bifurcation location depends on *κ*, however, the three mixtures are not equally distant from this boundary at fixed values of *R*_E+_ and *R*_*j*_. The bifurcation occurs at smaller *R*_E+_ for smaller *κ*; therefore, under the parameters of Fig. 3, low-*κ* networks lie farther inside the multistable regime and switch less frequently. The boundary regime is particularly challenging for mean-field predictions, as small changes in *R*_*j*_ can strongly alter branch stability and switching behavior, consistent with operation near a bifurcation. Across 45 independent connectivity realizations, switching behavior varied substantially, and some realizations showed no detected transition during the full 900 s observation window. Nevertheless, switching rates increased strongly with *κ*, whereas the number of non-switching realizations decreased (Supplementary Table 1). Comparison with the second parameter set in Supplementary Table 1 further shows that a small reduction in *R*_*j*_ substantially alters switching statistics, consistent with strong sensitivity to the distance from the bifurcation.

**Table 1.** Network architecture and clustering parameters, shared across mean-field, binary, and spiking implementations.

| Symbol / name | Value | Description |
| --- | --- | --- |
| $N_E$ | 8,000 | number of excitatory neurons |
| $N_I$ | 2,000 | number of inhibitory neurons |
| $N_Q$ | 20 | number of clusters |
| $n_E = N_E/N$ | 0.8 | excitatory fraction |
| $n_I = N_I/N$ | 0.2 | inhibitory fraction |
| $p_{EE}$ | 0.3 | baseline connection probability E→E |
| $p_{EI}$ | 0.3 | baseline connection probability I→E |
| $p_{IE}$ | 0.3 | baseline connection probability E→I |
| $p_{II}$ | 0.3 | baseline connection probability I→I |
| $g$ | 1.2 | inhibition strength parameter (BRN) |
| $R_{E+}$ | *varied* | excitatory clustering factor |
| $R_j$ | *varied* | relative inhibitory clustering, Eq. (R<sub>I+</sub>) |
| $\kappa$ | *varied* | clustering mix (0: probability, 1: weight) |
| Connectivity rule | Poisson multapses (main text) | $K_{ij} \sim \text{Poisson}(p_{ij})$ |

**Figure 4.**
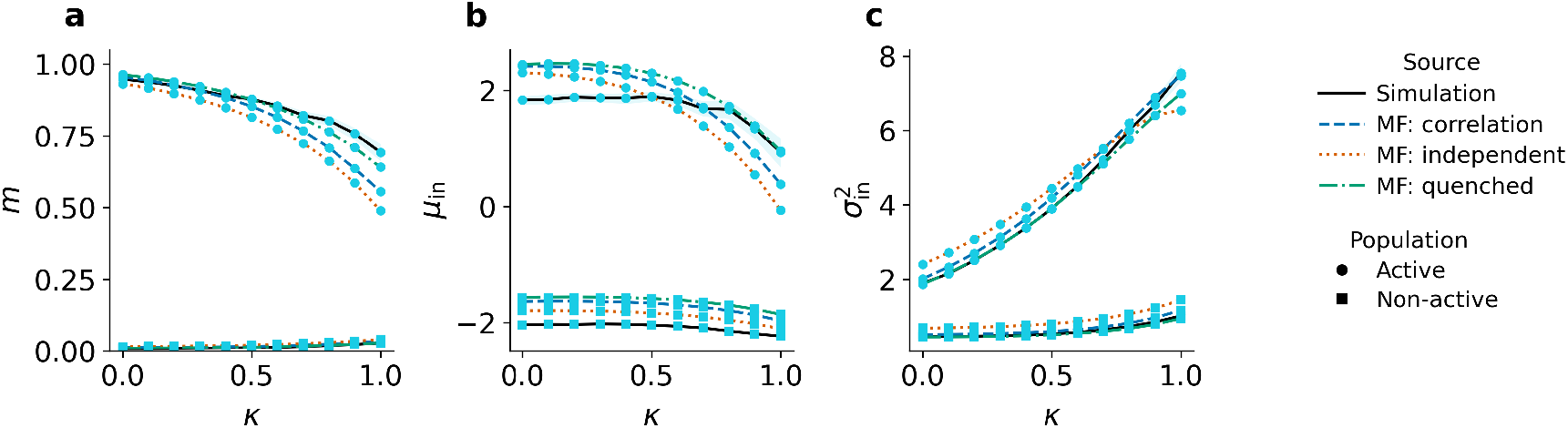
Quenched input variability captures the qualitative *κ*-dependence of single-active-cluster states. (a) mean activity *m*, (b) mean recurrent input *µ*_in_, and (c) recurrent-input variance 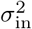 as a function of *κ*. Binary networks (*N*_*E*_ = 8,000, *N*_*I*_ = 2,000, *N*_*Q*_ = 20, *R*_E+_ = 7.25, *R*_*j*_ = 0.75) were simulated for 45 independent connectivity realizations and three initial conditions per realization. Each run was simulated for 900 s of model time after a warm-up period of 4 × 10^5^ asynchronous updates, with population activity sampled every 1.2 × 10^4^ updates, yielding 90,000 samples per run. Network states were inferred from the long activity traces of excitatory clusters using active-set state estimation with changepoint smoothing. Statistics were computed from all eligible single-active-cluster episodes. Repeated occurrences of the same state were first averaged within each run, followed by averaging across active-cluster identities, initial conditions, and connectivity realizations. The number and duration of eligible episodes varied across *κ*; the corresponding switching and state-repertoire statistics are reported in Supplementary Table 1. Lines show simulations and predictions from the quenched, independent, and correlated mean-field closures, computed using damped Picard iteration. Marker shape distinguishes active and inactive cluster populations, and shaded areas indicate bootstrap 95% confidence intervals across connectivity realizations.

The comparison with the mean-field predictions further reveals realization-specific heterogeneity. In the simulations, the activity of a high-activity state depends on the identity of the active cluster (Fig. 3a2–c2 and a3–c3). This asymmetry reflects quenched variability in the sampled connectivity, through which clusters differ in their realized input statistics, including their mean indegrees. By contrast, the standard mean-field theory uses theoretical connectivity expectations and therefore predicts identical activity levels for states that differ only in the identity of the active cluster. We therefore additionally evaluated an empirical mean-field theory in which the theoretical connectivity moments were replaced by cluster- and projection-specific means and variances measured from the corresponding connectivity realization. This breaks the symmetry between clusters and generally improves the prediction of high-activity states farther inside the multistable regime. At intermediate mixing (Fig. 3b), for example, states { 18 } and { 4 } are captured more closely than state { 17 } (Fig. 3b2–3). Closer to the bifurcation, particularly for weight clustering (Fig. 3c2,c3), both theories remain quantitatively less reliable and may fail to place the high-activity branch at the simulated parameter value. Realization-specific connectivity statistics thus explain part, but not all, of the discrepancy. To assess whether the displayed states are representative, we initialized the fixed-point solvers with state-resolved cluster activities extracted from longer simulations covering a broad range of active-cluster configurations (see Methods and Supplementary Fig. S3). Near the bifurcation, the iteration frequently converged to a different solution, including the uniform low-activity state. However, the returned solutions generally agreed more closely with another state observed in the corresponding simulation than with the state used for initialization. Thus, the predicted fixed points typically have counterparts in the simulated dynamics, but individual branches cannot always be recovered reliably. Remaining quantitative deviations likely reflect uncertainty in state-activity estimates and finite-size temporal fluctuations absent from the theory.

### Quenched input variability shapes cluster-state activity across *κ*

The residual discrepancies near the bifurcation, particularly for weight clustering in Fig. 3c and Supplementary Fig. S3, suggest that temporal fluctuations not represented by the quenched closure may influence the predicted fixed-point landscape. We therefore considered an independent closure, in which the variance of binary neuronal fluctuations is determined self-consistently by the mean activities, and a correlated closure adapted from covariance-aware mean-field approaches [54–56]. The latter approximates the recurrent input as a correlated Gaussian field and jointly determines mean activities and pairwise covariances, allowing correlations to affect the predicted fixed points (see Methods).

To compare these closures systematically outside the strongly realization-sensitive boundary regime, we reduced the relative inhibitory clustering from *R*_*j*_ = 0.79 to *R*_*j*_ = 0.75, moving the networks farther inside the multistable regime. Switching was substantially sparser, allowing single-active-cluster states to be identified more reliably. For each value of *κ*, we simulated 45 independent connectivity realizations, using three initial conditions per realization and averaged the state-resolved statistics across single-active-cluster states, initial conditions, and realizations. The frequency and duration of eligible episodes varied across *κ*; the corresponding switching and state-repertoire statistics are reported in Supplementary Table 1.

The mean activities of both active and inactive clusters vary systematically with *κ* (Fig. 4a), accompanied by corresponding changes in the mean and variance of the recurrent input (Fig. 4b,c). This dependence is qualitatively reproduced by the mean-field predictions. By construction, changing *κ* preserves the first moment of effective recurrent coupling for fixed population activities. Setting all input-variability terms to zero therefore eliminates the explicit *κ*-dependence of the predicted fixed points. Within the present interpolation, the *κ*-dependence of cluster activity is thus entirely mediated by input variability. Because probability and weight contrasts contribute differently to the second input moments, this variability necessarily changes when moving across the wiring–weight continuum, despite the matched first moment.

The hierarchy of closures identifies the dominant contribution to this effect. The quenched closure alone reproduces the qualitative dependence of active- and inactive-cluster activities on *κ*, whereas adding independent neuronal fluctuations or pairwise covariances modifies the predictions only modestly and does not consistently improve agreement. State-resolved comparisons further show that the correlated closure does not consistently reproduce the measured input and neuronal-state correlations in the activity regimes considered here (Supplementary Fig. S4). For fixed indegree connectivity, where the contribution of indegree variability to the quenched input variance is absent, the quenched closure does not recover the clustered fixed points, whereas the independent closure does (Supplementary Fig. S2). Thus, ensemble-level quenched input variance is the main contribution shaping the *κ*-dependent state activities in the Poisson networks, while independent fluctuations are sufficient to recover clustered fixed points in the fixed indegree theory. Pairwise correlations have comparatively little influence on the systematic *κ*-dependence in this regime, although small fluctuation corrections can become amplified near a bifurcation and alter branch stability. The residual discrepancies in this sensitive regime may therefore involve higher-order or non-Gaussian fluctuations, realization-specific connectivity heterogeneity, or difficulties in solver convergence, including possible jumps between different solution branches.

We additionally quantified pairwise correlations of recurrent input fields and neuronal states across the complete simulated trajectories, without conditioning on inferred network states (Fig. 5). Within-cluster input correlations decrease systematically with *κ*, consistent with a reduction in shared-input structure as cluster contrast is shifted from connection probabilities to synaptic weights. Across-cluster input correlations are substantially smaller but show a similar decreasing trend. Within-cluster state correlations are of comparable magnitude but vary only weakly with *κ*, while across-cluster state correlations remain negligible. Both input and state correlations depend on the average connection probability 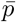. Thus, pairwise input correlations change systematically across the continuum, but neither the measured state correlations nor their inclusion in the mean-field closure substantially alters the principal *κ*-dependence of the single-cluster states.

**Figure 5.**
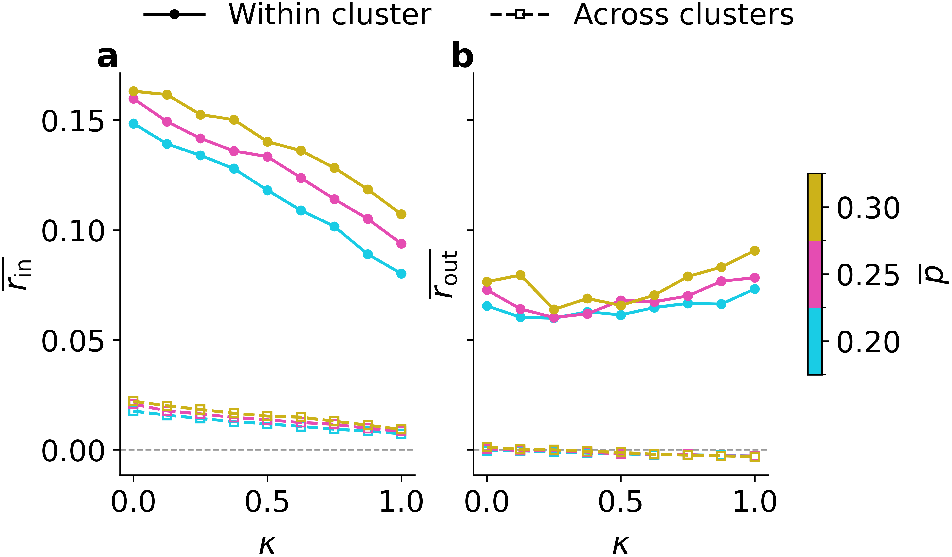
Input correlations decrease with κ, whereas neuronal-state correlations vary weakly. Binary networks (N_E_ = 8,000, N_I_ = 2,000, N_Q_ = 20, R_E+_ = 7.25, R_j_ = 0.79) were simulated for 45 independent connectivity realizations and three initial conditions per realization. After a warm-up period of 2×10^5^ asynchronous updates, correlations were estimated from 1.2×10^7^ updates by sampling every 1.2 × 10^4^ th update, corresponding on average to one full network update cycle, yielding 1,000 sampled configurations per run, corresponding to 10 s of model time. Correlations were computed over the complete sampled trajectories without conditioning on inferred network states. Pairwise Pearson correlations were calculated for sampled excitatory neuron pairs, Fisher-z transformed, averaged across pairs, initial conditions, and connectivity realizations, and transformed back to correlation coefficients. Mean correlations are shown as a function of κ for within-cluster (solid) and across-cluster (dashed) pairs and for different average connection probabilities 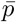 (color). (a) Correlations of recurrent input fields h_i_ . (b) Correlations of neuronal states σ_i_.

### Spiking LIF implementation preserves metastability across *κ* in simulation and on neuromorphic hardware

We test whether the binary network results transfer to a more biophysical setting by instantiating the same clustered architecture as a spiking LIF network (Fig. 6). For three tested mixtures *κ* ∈ { 0, 0.5, 1 }, activity exhibits metastable switching between cluster-activity states. Compared to the binary implementation, the increase in active-cluster rates toward smaller *κ* is less pronounced and does not require adaptation of *R*_*j*_.

**Figure 6.**
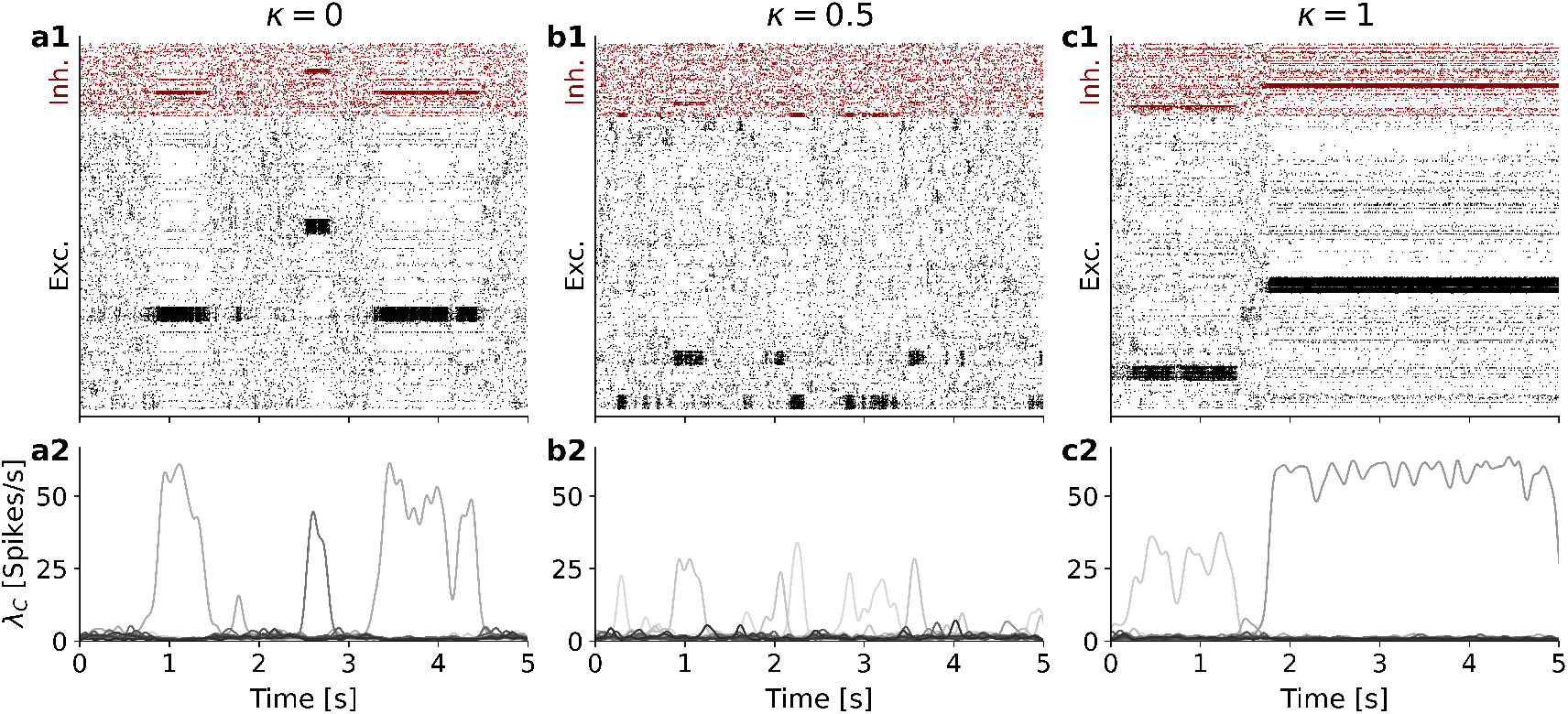
Spiking LIF networks reproduce metastable switching across clustering mixtures. Networks of N_E_ = 8,000 excitatory and N_I_ = 2,000 inhibitory leaky integrate-and-fire neurons were simulated with fixed parameters of excitatory clustering strength R_E+_ = 5 for N_Q_ = 20 clusters and relative inhibitory clustering strength R_j_ = 0.78 for (a) structural clustering (κ = 0), (b) mixed clustering (κ = 0.5), and (c) weight clustering (κ = 1). Top panels (a1–c1) show raster diagrams where each vertical tick represents the occurrence of an action potential. All three clustering regimes exhibit metastable switching, consistent with the binary network dynamics. Bottom panels (a2–c2) show per-cluster mean firing rates λc, computed by Gaussian kernel convolution of spike trains (σ = 50 ms, truncated at ±3σ). Connectivity fluctuations across random seeds can affect regime stability. A fixed indegree example is provided in the Supplementary Information.

Notably, robust metastability is obtained at a lower excitatory clustering strength *R*_E+_ than in the binary simulations. Together, these results show that metastable dynamics persist in structural, mixed, and weight-clustered regimes in both the abstract binary model and a spiking implementation.

We additionally implemented the clustered network architecture for the same three tested mixtures on the BrainScaleS-2 neuromorphic hardware platform [57] (Fig. 7). Due to hardware fan-in limits and the relatively high baseline connection probability, we used fixed indegree rather than Poisson connectivity, thereby eliminating realization-to-realization indegree variability while retaining disorder in presynaptic identities. The neuromorphic platform allowed calibration of neuron parameters to the target values. Nevertheless, as visible in Fig. 7, neurons remained heterogeneous: some neurons fired strongly, whereas others barely participated in the network dynamics. This variability may arise from the sampled connectivity or from residual uncertainty in the calibrated neuron parameters. All three networks produced the desired metastability despite neuron heterogeneity and the small population sizes of only 40 excitatory and 10 inhibitory neurons per cluster. Repeating the experiment 25 times with different connectivity seeds and neuron-to-cluster assignments robustly generated metastability as observed in each of the 25 emulations.

**Figure 7.**
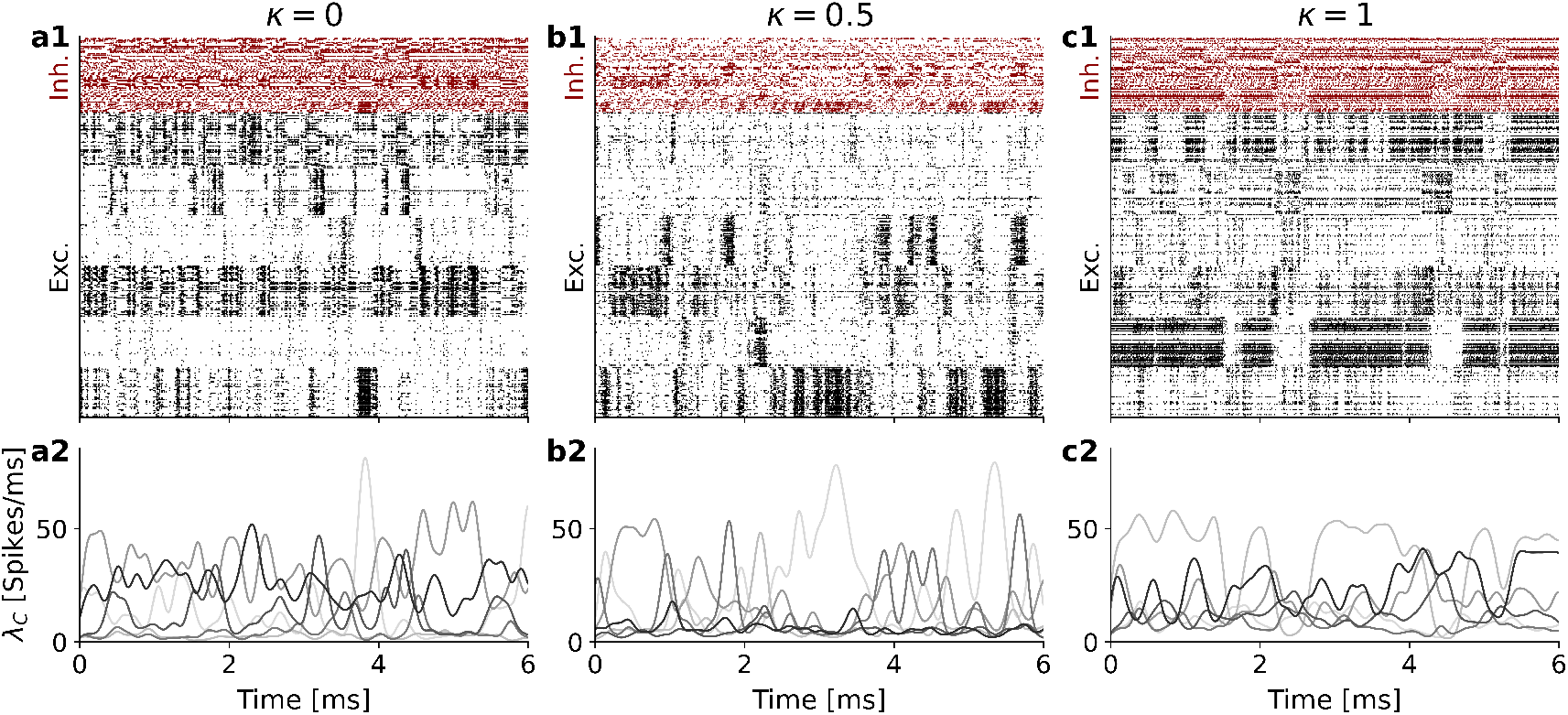
Neuromorphic implementation on BrainScaleS-2 exhibits metastable switching in small-scale networks. **Networks** of N_E_ = 240 excitatory and N_I_ = 60 inhibitory leaky integrate-and-fire neurons across N_Q_ = 6 clusters were emulated on BrainScaleS-2 with fixed excitatory clustering strength R_E+_ = 4.5 and relative inhibitory clustering strength R_j_ = 0.65 for (a) structural clustering (κ = 0), (b) mixed clustering (κ = 0.5), and (c) weight clustering (κ = 1). To avoid high fan-in from random network construction, fixed indegree connectivity was used with baseline connection probability p = 0.15. Top panels (a1–c1) show raster diagrams; each vertical tick indicates an action potential. All three clustering regimes exhibit metastable switching. Bottom panels (a2–c2) show excitatory per-cluster firing rates λc, computed by Gaussian kernel convolution of spike trains (σ = 100 µs, truncated at ±3σ). The displayed time window shows the first 6 ms of each emulation.

### Discussion

### What connectivity means biologically, experimentally, and in the model

In the central nervous system of both vertebrates and invertebrates, a presynaptic neuron commonly forms multiple synaptic contacts onto the same postsynaptic partner [58–62], including cell-type-dependent autapses [63–66]. At this detailed level of anatomical description, connectivity is therefore naturally non-binary. A multigraph, in which a neuron pair can be linked by multiple edges, therefore provides a natural generalization representing pairwise connectivity in terms of synapse count. This also allows for functional heterogeneity because multiple contacts increase the range of effective connection strengths, provide redundancy against synaptic transmission failures at individual contact sites [67], and introduce independent variability of vesicle release across contact sites [68].

The multigraph is only one possible abstraction. Empirical distributions of synapse counts can deviate substantially from a simple Poisson model, making it useful to separate two questions: whether a connection exists at all (≥1 synapse), and how many synaptic contacts are formed conditional on connection. Compared to the multigraph, such a two-stage definition could reduce the number of distinct presynaptic partners while increasing multiplicity among existing partners, thereby concentrating input onto fewer sources and altering input overlap.

Within this broader heterogeneity, our model isolates one effective degree of freedom: for a fixed contrast between within- and across-cluster interactions, the contrast can be implemented through contact statistics, synaptic efficacy, or mixtures of both. The mixing parameter *κ* is therefore not intended to correspond one-to-one to a single biological variable, but provides a mesoscale coordinate for comparing circuit implementations. Its structural component relates to connection probability, contact multiplicity, and local versus long-range projection statistics obtained from anatomical or connectomic data, whereas its weight component relates to synaptic efficacy and efficacy distributions estimated from physiological measurements. Intermediate *κ* values represent cases in which anatomical organization and synaptic-strength heterogeneity jointly shape effective clustered coupling. Because shifting contrast between connection probabilities and weights changes input variance, shared-input statistics, and fixed-point structure even when mean input is preserved, comparable assembly dynamics need not arise from identical microscopic circuit parameters. In the present formulation, synaptic efficacies are fixed within each projection class; replacing them with empirically motivated efficacy distributions would be a natural extension.

This interpretation is relevant for comparisons across brain areas or species, where recurrent organization may differ not only in connection density but also in synapse number, projection range, and efficacy distributions. In such settings, *κ* provides a phenomenological way to ask whether similar assembly-like dynamics could be supported by different combinations of anatomical and physiological constraints. Intermediate *κ* values are particularly useful conceptually, because future datasets combining dense connectomics with large-scale physiological measurements may constrain both contact statistics and synaptic-strength distributions rather than either quantity alone. In the present model, synaptic efficacies are fixed within each projection class; replacing these values by empirically motivated efficacy distributions would be a natural extension.

A separate challenge is to infer these structural and physiological components from empirical data. Reported connectivities can refer to different underlying quantities. Importantly, only physiological measurements such as intracellular recordings of synaptic currents can provide reliable estimates of effective connection strength, while structural (anatomical and morphological) data are required to reconstruct pairwise connection patterns, or even synapse- or bouton-level counts of polyadic contact points at the ultrastructure level in electron microscopy data. Thus, even when non-random local connectivity is detected, it may remain difficult to determine whether an apparent cluster structure is implemented primarily through structure (*p*-contrast) or synaptic efficacy (*J* -contrast), especially under partial observation. Small samples can make distinct generative mechanisms appear similar, and signatures often attributed to clustering may also arise from alternative forms of organization such as distance dependence or broad degree distributions [69]. This ambiguity persists in large-scale recordings, where connectivity inferred from extracellular activity remains sensitive to unobserved neurons, common input, and sampling biases [70–72].

### Embedding, scale, and plasticity as determinants of effective clustering

Clustered network models can link metastable dynamics to behavior in animal experiments [23, 42, 73]. However, motif-level models necessarily abstract away the fact that, in cortex, assembly-like dynamics unfold within broader recurrent circuitry and are likely influenced by ongoing inputs and global state. Moreover, if clustered structure is present, it need not appear as a “pure” motif but may be expressed as an effective organization intermingled with other circuit features [69]. In line with the common experimental and theoretical terminology, we interpret assemblies as functional population states rather than isolated modules: neurons can contribute to multiple population patterns over time [6, 25, 74–76], and the ensembles expressed can vary with context, task demands, and global state [77]. From this perspective, external inputs are not merely background drive for balance; they can bias which states are stabilized and how transitions unfold by altering effective population-level interactions [21, 78]. Our model does not include such embedding or input-dependent reconfiguration, so how clustered motifs interface with other circuit features to produce richer, task-specific dynamics remains an open research question.

Interpretations of clusters depend on scale. At the scale of small assemblies, clusters can be defined operationally at the population level as recurring co-activation patterns. At larger scales, clustered structure may reflect additional organization, including hierarchical or multi-level motifs [35]. Moreover, in large-scale and multi-area circuit models, structured recurrent connectivity can give rise to slow, state-like population dynamics that are often described as metastable or metastable-like [79, 80]. This motivates the distinction between local and global attractor-like states, and viewing clustering as one level within a broader hierarchy of recurrent organization.

Plasticity provides a plausible route to intermediate mixtures, but the mapping from biological mechanisms to our parameterization is best interpreted at the level of effective connectivity rather than as a one-to-one correspondence with a single process. In models where clustered or attractor-like structure emerges primarily through changes in synaptic efficacy driven by local learning rules, the resulting organization is naturally closer to the weight-dominated end of our continuum (larger *κ*) [43,44,81]. Conversely, contact-level structural plasticity provides a plausible mechanism by which experience can bias effective connectivity and increase effective within-group coupling in models, consistent with a shift toward smaller *κ* in our phenomenological continuum [61, 82]. At the same time, spine dynamics and turnover complicate a direct mapping from anatomical contacts to functional coupling, so changes in contact statistics need not translate one-to-one into circuit-relevant connectivity [83]. Mixed cases, in which clustered structure is partly imposed by wiring constraints and partly refined or stabilized by synaptic plasticity, conceptually align with intermediate *κ* values. Such mixed structural–synaptic mechanisms may also help stabilize learned structure: Hebbian plasticity alone is typically unstable [84], so maintaining persistent representations and preventing runaway dynamics likely requires compensatory processes such as synaptic scaling and inhibitory plasticity [43, 85–88]. Moreover, presynaptic connectivity structure can shape how plasticity sculpts excitation and inhibition, influencing the emergence of EI co-tuning [89]. Relatedly, stable task performance can coexist with ongoing representational drift of single neurons, implying that stability may be expressed at the level of population codes rather than fixed membership of individual assemblies [90, 91].

### Neuromorphic implementation illustrates trade-offs between synapse count and weight resolution

Attractor networks are computational primitives commonly used in neuromorphic computing [48, 92]. Soft winner-take-all (sWTA) motifs have been used in applications such as state machines, contrast enhancement, and decision making [15, 93–98]. Compared with WTA and persistent-activity attractor motifs, neuromorphic demonstrations that explicitly foreground metastable switching as the primary computational primitive remain comparatively rare but are increasingly explored both as efficient substrates for neuroscience-oriented models and as bio-inspired computational architectures that exploit stochastic switching constructively, for example in generative sampling with suitable learning rules [99], reinforcement-learning style mechanisms [100], and stochastic search [101]. Their hardware implementation remains challenging, because switching statistics can be sensitive to noise, finite-size effects, and device mismatch, making predictable transition rates across chips and operating conditions difficult to achieve without calibration or adaptive control [102, 103]. Robust network design can help mitigate these challenges. For example, EI clustering can support metastability over a broad parameter regime [36, 104], although our framework is not restricted to this case and also applies to E-clustered and other clustered architectures. At the same time, neuromorphic substrates impose practical constraints on routing and fan-in/fan-out, delays, network size, and synaptic weight resolution [48, 49, 105]. As a result, network mappings must respect available routing resources, synapse formats, and discretized, limited-range weights [48,57,106,107]. In this context, combining weight-based and connectivity-based structure can help adapt clustered network models to such hardware constraints.

In hardware terms, *κ* specifies where the cluster contrast is primarily represented. At *κ* = 0, all synaptic weights remain at their projection-specific baseline values, so the network requires only the four baseline weight classes *J*_EE_, *J*_EI_, *J*_IE_, *J*_II_, while cluster contrast is carried by the routing pattern and contact multiplicity. For any *κ >* 0, within- and across-cluster weights differ, yielding up to eight projection- and cluster-relation-specific weight classes 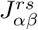 (Eq. (3)) and shifting implementation demands toward weight precision, dynamic range, and calibration. However, the present interpolation also changes the expected indegree: although the effective mean input is preserved, the probability modulation has an average factor *ρ*_*K*_(*κ*; *R, N*_*Q*_) that equals one at the endpoints *κ* = 0 and *κ* = 1, but can fall below one for intermediate *κ*. For strong cluster contrasts, this reduction can amount to several tens of percent. Intermediate *κ* values can reduce the expected number of realized contacts, while both intermediate and wiring-dominated implementations can reduce the expected number of distinct presynaptic partners (fan-in). Intermediate *κ* values can therefore provide a hardware-relevant compromise by reducing the routed synapse count on substrates that represent individual contacts separately, while requiring a smaller weight contrast than pure weight clustering. This potential advantage is substrate-dependent, because changes in contact count and indegree also alter input variance and may affect the robustness of the metastable regime. After hardware quantization, very small nonzero values of *κ* may also collapse to fewer effective weight levels. Thus, *κ* can be interpreted as a compilation parameter for adapting the same clustered dynamical motif to substrates with different routing, fan-in, weight-resolution, and calibration constraints.

The multigraph realization further highlights this hardware dependence. If a substrate can merge multapses into a single synapse with an integer-multiple or otherwise scaled effective weight, contact multiplicity can be represented compactly. If multapses must be represented as separate synaptic resources, the wiring-dominated regime increases memory and routing demands while keeping the nominal weight set simple. Thus, neither endpoint is universally preferable; the useful operating point depends on the available routing resources, fan-in/fan-out limits, synapse format, weight resolution, and calibration precision of the target substrate. Early neuromorphic demonstrations of attractor-like dynamics and long-lived bistable states suggest feasibility of hardware implementations, but scaling of networks, network calibration under mismatch, and achieving predictable switching statistics remain open engineering challenges [101, 103, 108].

### Limitations of the mean-field analysis

Our mean-field analysis is approximate: it provides qualitative stability guidance, but it omits higher-order correlations and finite-size fluctuations that set metastable dwell times and switching rates in finite networks [22, 109], especially near stability boundaries [110]. Moreover, while *κ* is meant to shift coupling contrast between connection probabilities and synaptic weights, in a directed Poisson multigraph this shift also changes contact-count variability and overlap statistics, thereby affecting higher input moments. Employing a hierarchy of mean-field closures taking into account different types of second-order variability, we show that *κ*-related changes in the quenched variability of the Poisson graph are the main driver behind changes in fixed point statistics. This sensitivity matters both biologically and for neuromorphic mapping, where robustness to structural variability and device mismatch is essential. To reduce sensitivity to connectivity disorder, we introduce a fixed indegree construction in the Supplementary Information. In this case, independent temporal variability and its dependence on *κ* are the dominant determinants for the fixed-point behavior. Beyond mean-field, higher-order-correlation- and finite-size-aware theories will improve quantitative predictions. Additional simplifications include homogeneous neuron parameters and limited robustness sweeps over network size, cluster count, sparsity, and external drive; our simulations therefore emphasize qualitative regime transfer rather than invariance of all *κ*-dependent signatures.

### Conclusion

Implementing clustered coupling via a contrast in connection probability (*κ* = 0) versus synaptic weight (*κ* = 1) defines a practical translation axis rather than a pure reparameterization. Although the expected mean recurrent input is preserved, varying *κ* changes expected indegree and higher input moments, thereby reshaping the fixed-point structure, state activities, input correlations, and the balance between single- and multi-cluster episodes. In the Poisson networks, ensemble-level quenched input variance accounts for the principal systematic dependence of cluster-state activities on *κ*, while realization-specific heterogeneity of connectivity becomes particularly important near bifurcations. Comparable metastable regimes can therefore occur at different absolute clustering strengths and may require retuning when transferred across substrates. Near bifurcation boundaries, transferring parameters without retuning can alter or abolish metastability. Mechanistically, we attribute these changes to altered input overlap and effective distance to stability boundaries, which modulate dwell statistics and switching. Binary, spiking, and neuromorphic implementations support metastable switching across the structural, mixed, and weight-clustering regimes, although their quantitative operating regimes and switching statistics differ. Together, these results establish *κ* as a useful design coordinate for transferring clustered metastable dynamics across software and neuromorphic substrates with different wiring and weight constraints.

## Methods

### Binary network simulations

We simulate asynchronous binary dynamics with single-neuron updates [36]. At update step *k* a neuron *i* is drawn, and its state *σ*_*i*_ ∈ {0, 1} is set to

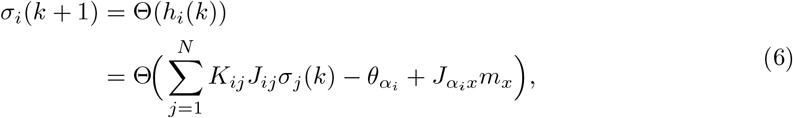

while all other neurons keep their state. Here *m*_*x*_ is a constant external drive coupled with strength *J*_*αx*_, and *K*_*ij*_ is drawn from the Poisson multigraph model. Baseline weights are 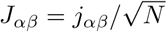, and clustered modulation follows Eq. (3). In the binary simulations, the external drive couplings were fixed as

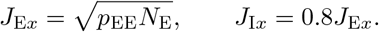

To allow heterogeneous update rates, each neuron *i* is assigned an update time constant *τ*_*i*_. At each simulation step, one neuron is sampled for update with probability

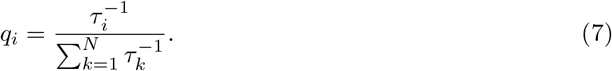

If *τ*_*i*_ = *τ*_*αi*_ depends only on neuronal type, then

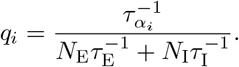

Time is reported in physical units by assigning a physical update interval to the expected number of simulation steps between consecutive updates of the same neuron. This convention is chosen for interpretability; it affects only the mapping between simulation steps and physical time, which depends on network size, and does not alter the network dynamics. Unless stated otherwise, we used the parameters in Table 1 for the main-text simulations and the parameters in Table 2 for the mean-field analysis and binary-network simulations.

**Table 2.** Binary / mean-field dynamics parameters.

| Parameter | Value | Description |
| --- | --- | --- |
| $\tau_E$ | 10 ms | excitatory update constant |
| $\tau_I$ | 5 ms | inhibitory update constant |
| $\theta_E$ | 1.0 | excitatory threshold |
| $\theta_I$ | 1.0 | inhibitory threshold |
| $m_x$ | 0.03 | constant external drive (rate) |

### Mean-field analysis

We describe the clustered binary network at the level of populations defined by type and cluster. Let *m*_*αr*_(*t*) ∈ [0, 1] denote the mean activity of type *α* ∈ {E, I} in cluster *r* ∈ {1, …, *N*_*Q*_}. Under a Gaussian approximation to the input distribution [55, 111], the population activities obey

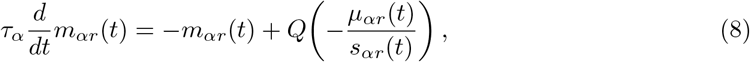

Where

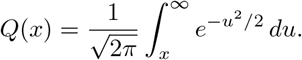

The *Q*-function gives the probability that a Gaussian input variable with mean *µ*_*αr*_ and standard deviation *s*_*αr*_ exceeds threshold. Thus, in the quasi-static limit,

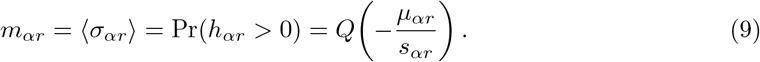

Because clustering enters only through type- and cluster-dependent modulations, we use block-constant effective parameters

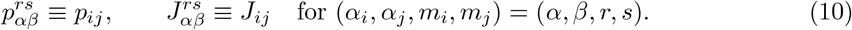

For equal-size clusters, 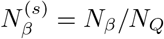. With constant external drive *m*_*x*_ and coupling *J*_*αx*_, the mean input is

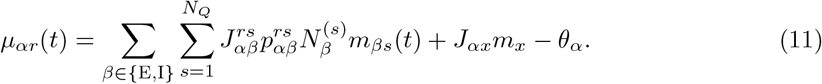

We obtain two covariance-free approximations: quenched and independent. For Poisson multapses *K*_*ij*_ ∼ Poisson(*p*_*ij*_) and presynaptic binary states with mean *m*_*βs*_(*t*), the variance contribution of one presynaptic neuron is

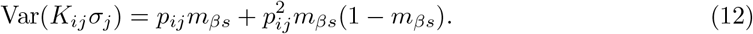

In the quenched approximation, we retain only the first term, which reflects variability due to the Poisson multapse statistics:

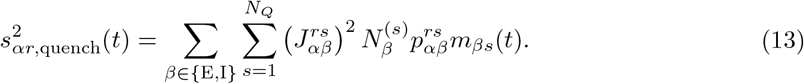

In the independent approximation, we additionally include the variance of the presynaptic binary state, assuming independent Bernoulli activity. For binary variables, this independent activity variance is fixed by the mean,

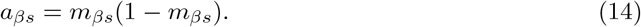

The corresponding input variance is

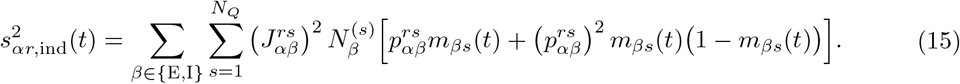

Thus, the quenched approximation includes only connectivity-induced variance, whereas the independent approximation includes both connectivity-induced variance and independent presynaptic activity variance. Both approximations neglect pairwise covariances between presynaptic units; these are included only in the correlation-sensitive extension below.

The quenched approximation was used for the analyses in Fig. 2 and Fig. 3. The independent approximation was introduced in Fig. 4 and Supplementary Fig. S3 and S4. Both approximations are self-consistent at the level of mean population activities, but they neglect pairwise covariances and may therefore underestimate fluctuations in regimes where collective variables induce correlations.

To estimate covariance contributions, we also used a correlation-sensitive extension [54–56]. In this formulation, the full input variance is

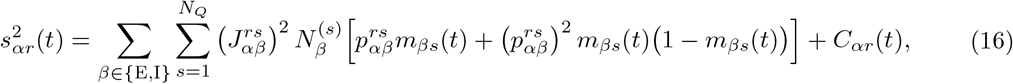

where *C*_*αr*_(*t*) collects covariance contributions between presynaptic populations:

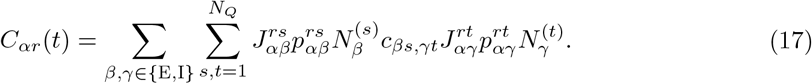

Here *c*_*βs,γt*_ denotes the averaged pairwise cross-covariance between blocks. The covariances are obtained from the self-consistency equation

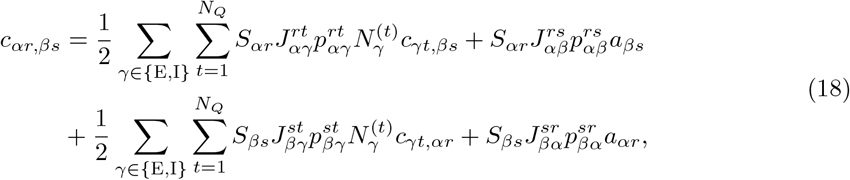

with susceptibility

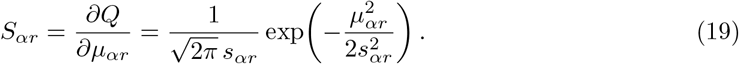

The correlation-sensitive formulation therefore augments the activity fixed-point problem by the block-pair covariances *c*_*αr,βs*_.

For constant input, fixed points satisfy

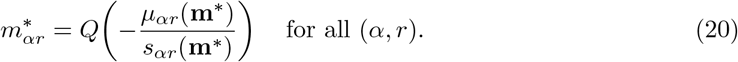

We located solutions using an effective response function approach [53], in which the 2*N*_*Q*_ population activities are reduced by symmetry to four macroscopic variables,

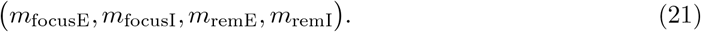

For a prescribed *m*_focusE_, we solved the remaining self-consistency equations and computed the implied output activity of the focus group. Fixed points correspond to intersections between prescribed input activity and implied output activity. Local stability was assessed from the Jacobian of Eq. (8) evaluated at **m**^∗^; a fixed point was classified as stable when all eigenvalues had strictly negative real parts.

Numerically, fixed points were obtained by solving

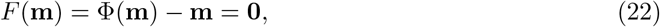

where Φ denotes the Gaussian mean-field response defined above. We used the SciPy hybrid solver, `hybr`, initialized from prescribed population activity vectors. The root tolerance was 10^−9^, with at most 4000 function evaluations. Solutions were accepted when the solver converged or when the maximum residual norm was below 5 × 10^−4^.

As a complementary numerical check, we also used damped Picard iteration,

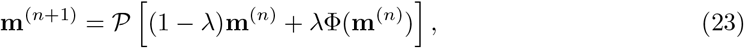

where *P* projects activities to the admissible interval [0, 1]. Unless stated otherwise, the damping parameter was *λ* = 0.1, the convergence tolerance was 10^−14^, and the maximum number of iterations was 10^5^. Convergence was assessed from the projected fixed-point residual.

For the correlation-sensitive theory, covariances were solved self-consistently together with the mean activities. For a given activity vector, the covariance update *C* 1→ Ψ(**m**, *C*) was iterated as

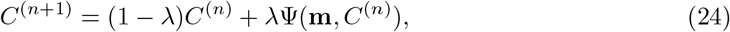

where *λ*Ψ(**m**, *C*^(*n*)^) is the right-hand side of Eq. (18). We then enforced numerical symmetry of *C*(^*n*+1^) via

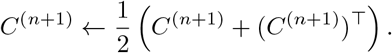

The same damping, tolerance, and maximum iteration count were used unless stated otherwise. The search space for fixed points is therefore increased from 2*N*_*Q*_ variables to 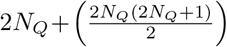 variables.

### Spiking network implementation

To test robustness in a more biophysical setting, we instantiate the same clustered architecture as a network of leaky integrate-and-fire (LIF) neurons with exponential synaptic currents, following [23]. Neuron *i* has type *α*_*i*_ ∈ {E, I} and evolves according to

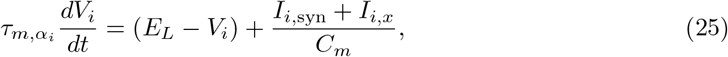

where *E*_*L*_ is the leak reversal, 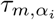the membrane time constant, *C*_*m*_ the capacitance, and *I*_*i,x*_ a constant external current. When *V*_*i*_ reaches threshold 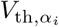, a spike is emitted, *V*_*i*_ is reset to *V*_*r*_ and held during an absolute refractory period *τ*_*r*_ (parameters in Table 3). We reuse the same abstract clustered architecture as in the binary network. In the main-text spiking simulations, connectivity is realized as a Poisson multigraph with synapse multiplicities *K*_*ij*_ ∼ Poisson(*p*_*ij*_), where *p*_*ij*_ follows the clustering rule in Eq. (3); the supplementary fixed indegree variant is described separately in the Supplementary Information. Baseline synaptic amplitudes follow the scale-free balanced calibration: *j*_*αβ*_ are given by Eq. (1) and mapped to weights via 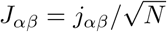, and clustered synapses use *J*_*ij*_ from Eq. (3). Equivalently, at the block level 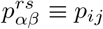 and 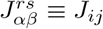 for (*α*_*i*_, *α*_*j*_, *m*_*i*_, *m*_*j*_) = (*α, β, r, s*). The synaptic current is the sum of excitatory and inhibitory components, *I*_*i*,syn_ = *I*_*i*,ex_ + *I*_*i*,in_, governed by

**Table 3.**
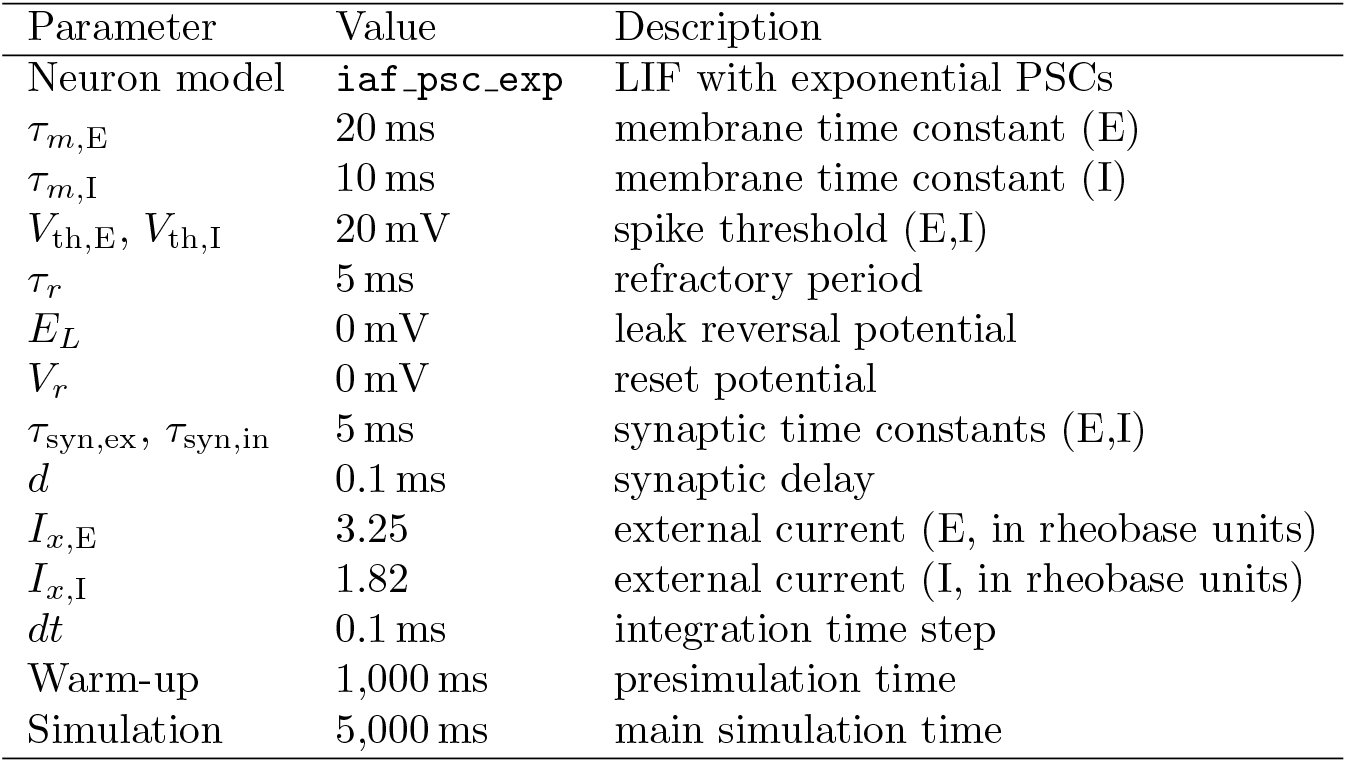
Spiking (NEST) neuron and simulation parameters.

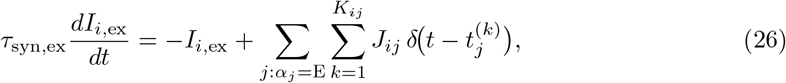

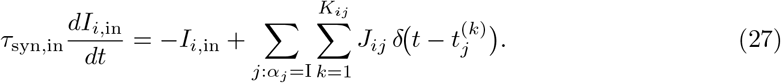

In the spiking model, the effective impact of a synaptic event depends on *τ*_*m*_ and 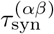 Following [23], we quantify this by the peak postsynaptic potential 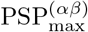 and define an effective coupling

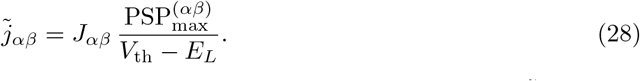

Imposing the same balanced-state conditions as in the binary network, but in terms of 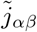, yields the spiking calibration reported in [23]. Both the binary and spiking implementations realize the same abstract clustered balanced network at the level of scale-free couplings *j*_*αβ*_ and the same block-structured multigraph connectivity. The binary mean-field analysis thus captures how clustered second-order input statistics shape population dynamics as an approximation to the spiking network, even though the transfer function from Gaussian input to output firing rates differs between binary threshold units and LIF neurons.

### Analysis of binary network simulations

Binary simulations store the initial state ***σ***(0) and, for each update step, the updated neuron index *i* and its state change Δ*σ*_*i*_ ∈ {−1, 0, +1}. Event rasters are generated from the ordered pairs (*i, t*(*k*)) associated with update steps *k* for which Δ*σ*_*i*_ = +1, corresponding to transitions into the active state. Population activities *m*_*αr*_(*t*) are reconstructed online by updating only the affected population,

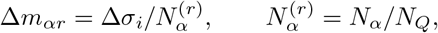

before temporal smoothing by convolution with a Gaussian kernel truncated to ± 3*σ*. For each network and initialization we compute Pearson correlations between the state ***σ*** and the recurrent input field **h** evaluated from the same sampled states, where

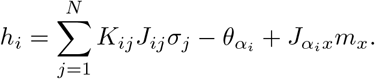

Correlations are estimated from random neuron pairs pooled over all sampled time points. Coefficients are Fisher-*z* transformed, averaged across pairs and initial conditions, and back-transformed for display. Error bars are obtained by bootstrap across connectivity realizations using per-network averaged Fisher-*z* values.

### State analysis

Network states were inferred from the sampled excitatory population activities using a multistage procedure comprising robust normalization, changepoint detection, active-set inference, and post-processing. Population activity was sampled every 10 ms during 900 s simulations. Piecewise-stationary activity intervals were identified using PELT [112] with a weighted (*ℓ*_2_) segment cost. Each interval was then assigned a binary active-set label by separating clusters into low- and high-activity components. Brief excursions and weak spillover activity were removed, and the resulting states were restricted to at most two active clusters. The final state sequence was used to calculate switching frequency, number of states, and state-specific population emissions. State inference used only excitatory activity, whereas emission estimates included both excitatory and inhibitory populations. Full details of the preprocessing, model fitting, parameter choices, and post-processing are provided in the Supplementary Information.

### BrainScaleS-2 implementation

We implemented a scaled version of the clustered spiking network on the BrainScaleS-2 neuromorphic system [57], using the PyNN `brainscales2` backend and the `iaf_psc_exp` neuron model. The network contained *N*_E_ = 240 excitatory and *N*_I_ = 60 inhibitory neurons arranged into *N*_*Q*_ = 6 clusters. Connectivity followed the same cluster-dependent construction as in the software simulations, but used fixed indegree to avoid excessively high fan-in on the hardware. We used baseline connection probability *p* = 0.15, excitatory clustering strength *R*_E+_ = 4.5, and relative inhibitory clustering strength *R*_*j*_ = 0.65. Three representative points along the mixing parameter *κ* ∈ { 0, 0.5 }, 1 were tested. Robustness was assessed by constructing 25 networks with different connectivity seeds and different neuron-to-cluster assignments. We screened raster plots of the initial 6 ms of each emulation to identify the presence of metastable switching. Due to the approximately 1,000-fold accelerated dynamics of BrainScaleS-2, membrane, synaptic, refractory, and delay time constants are given in hardware time. We used *τ*_*m*,E_ = 20 µs, *τ*_*m*,I_ = 10 µs, *τ*_syn,ex_ = *τ*_syn,in_ = 5 µs, refractory period 5 µs, and synaptic delay 0.1 µs. Each run consisted of an 80 ms warm-up followed by 100 ms of recorded activity. Neurons were calibrated using the BrainScaleS-2 software framework before execution. Experiments were executed on the BrainScaleS-2 accelerated mixed-signal neuromorphic platform accessed through the EBRAINS Research Infrastructure.

### AI usage

In parts of this work, we used the AI-based code assistant Codex (OpenAI) to generate and debug analysis and simulation code. To improve the clarity and readability of the manuscript, we used ChatGPT (OpenAI) to suggest alternative phrasings and to help streamline the text. All code and text suggestions were carefully reviewed, verified, and, where necessary, modified by the authors, who retain full responsibility for the final content.

## Code availability

All code necessary to reproduce the analyses and simulations presented in this study is available in the public GitHub repository: https://github.com/nawrotlab/EI-Clustering.

## Acknowledgements

We thank Thomas Rost for providing the code for the binary network simulations from [36]. Figure 1a: Created in BioRender https://BioRender.com/lefi1zt. This work was funded by the Deutsche Forschungsgemeinschaft (DFG, German Research Foundation) – Project-ID 431549029 – SFB 1451, project A06. We also thank the IT Center University of Cologne (ITCC) for providing compute resources and support on the DFG-funded HPC system RAMSES (Research Accelerator for Modeling and Simulation with Enhanced Security) (DFG funding number: INST 216/512-1 FUGG). The EBRAINS Research Infrastructure provided access to the BrainScaleS-2 neuromorphic computing system. We thank the organizers of the CapoCaccia Workshop Towards Neuromorphic Intelligence 2024, for enabling the neuromorphic implementation. We especially thank Yannik Stradmann and Sebastian Billaudelle for their help with the BrainScaleS-2 platform.

