## Supplementary Information for "Metastable Neural Assemblies on a Wiring–Weight Continuum"

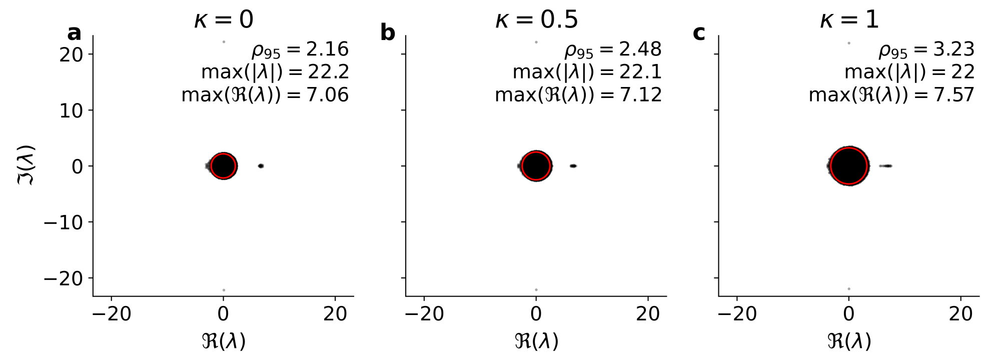


**Figure S1.** Eigenvalue spectrum of the multigraph connectivity matrix with Poisson-distributed synapse counts ($N_E = 8000$, $N_I = 2000$, $N_Q = 20$, $R_{\mathrm{E}+} = 7.25$, $R_j = 0.75$). The bulk radius $\rho_{95}$ is defined as the 95th percentile of $|\lambda|$ and is shown as a red circle. $\rho_{95}$ increases with $\kappa$, suggesting greater potential for linear amplification of activity fluctuations. In contrast, $\max|\lambda|$ remains approximately constant and likely reflects the block structure of the EI-clustered network. Note that this spectrum is computed for $W$ and should not be confused with the effective Jacobian, which depends on the operating point and thus differs for each network state. The complex-conjugate eigenvalue pair lying outside the bulk and giving the largest magnitude, $\max|\lambda|$, is associated with the global activation mode of the network due to the balanced condition of the network [3].

### Alternative multigraph realization: fixed indegree construction per block

The main text uses a Poisson multapse model $K_{ij} \sim \text{Poisson}(p_{ij})$. Here we describe an alternative construction that fixes the indegree per presynaptic block while allowing multapses and autapses. Importantly, pairwise multapse statistics remain close to Poisson in the sparse regime, so finite-size variance is not substantially reduced.

Neuron $i$ has type $\alpha_i \in \{\text{E}, \text{I}\}$ and cluster $m_i = r$. For presynaptic type $\beta$ and cluster $s$, define the presynaptic block

$$\mathcal{S}_{\beta s}(i) = \{j \mid \alpha_j = \beta,\ m_j = s\}, \qquad \lambda_{i,\beta s} = \sum_{j \in \mathcal{S}_{\beta s}(i)} p_{ij}.$$

Because $p_{ij}$ is block-constant for fixed $(\alpha_i, \beta, r, s)$, $\lambda_{i,\beta s}$ depends only on $(\alpha_i, r, \beta, s)$.

For each postsynaptic neuron $i$ and block $(\beta, s)$, we set a target indegree

$$K^{\text{tar}}_{i,\beta s} = \lfloor \lambda_{i,\beta s} \rceil, \tag{1}$$

and sample $K^{\text{tar}}_{i,\beta s}$ presynaptic partners with replacement from $\mathcal{S}_{\beta s}(i)$ with probabilities

$$\pi^{(\beta s)}_{ij} = \frac{p_{ij}}{\sum_{k \in \mathcal{S}_{\beta s}(i)} p_{ik}}, \qquad j \in \mathcal{S}_{\beta s}(i). \tag{2}$$

Let $K_{ij}$ be the number of times neuron $j$ is selected from its block. Then, for fixed $(i, \beta, s)$, the vector $\big(K_{ij}\big)_{j \in \mathcal{S}_{\beta s}(i)}$ has a multinomial distribution with total count $K^{\text{tar}}_{i,\beta s}$ and probabilities $\pi^{(\beta s)}_{ij}$.

The block indegree is deterministic,

$$K_{i,\beta s} = \sum_{j \in \mathcal{S}_{\beta s}(i)} K_{ij} = K^{\text{tar}}_{i,\beta s}, \qquad \text{Var}(K_{i,\beta s}) \approx 0,$$

up to $\mathcal{O}(1)$ rounding in (1). In contrast, the pairwise multiplicities satisfy

$$\mathbb{E}[K_{ij} \mid K^{\text{tar}}_{i,\beta s}] = K^{\text{tar}}_{i,\beta s}\, \pi^{(\beta s)}_{ij}, \tag{3}$$

$$\text{Var}(K_{ij} \mid K^{\text{tar}}_{i,\beta s}) = K^{\text{tar}}_{i,\beta s}\, \pi^{(\beta s)}_{ij}\big(1 - \pi^{(\beta s)}_{ij}\big), \tag{4}$$

$$\text{Cov}(K_{ij}, K_{ik} \mid K^{\text{tar}}_{i,\beta s}) = -K^{\text{tar}}_{i,\beta s}\, \pi^{(\beta s)}_{ij} \pi^{(\beta s)}_{ik} \qquad (j \neq k), \tag{5}$$

i.e., multinomial variance with negative within-block covariances. Since blocks are large, $\pi^{(\beta s)}_{ij} \ll 1$ (for uniform sampling, $\pi^{(\beta s)}_{ij} = 1/|\mathcal{S}_{\beta s}(i)|$), hence $1 - \pi^{(\beta s)}_{ij} \approx 1$ and

$$\text{Var}\big(K_{ij} \mid K^{\text{tar}}_{i,\beta s}\big) = K^{\text{tar}}_{i,\beta s}\, \pi^{(\beta s)}_{ij}\big(1 - \pi^{(\beta s)}_{ij}\big) \approx K^{\text{tar}}_{i,\beta s}\, \pi^{(\beta s)}_{ij} \approx p_{ij}.$$

This equals the leading-order variance in the Poisson multapse model ($\text{Var}(K_{ij}) = p_{ij}$). Therefore, fixing the block totals $K_{i,\beta s}$ removes fluctuations of the *total* indegree per block, but does not substantially reduce pairwise multapse variance in the sparse regime considered here. In finite networks the two constructions can nevertheless behave differently. A likely contributor is that Poisson multapses induce block-level indegree fluctuations $\text{Var}(K_{i,\beta s}) = \lambda_{i,\beta s}$, whereas the fixed-block scheme suppresses this quenched heterogeneity, reducing the occurrence of strongly driven neurons with unusually large input that can nucleate cluster activation via recurrent gain.

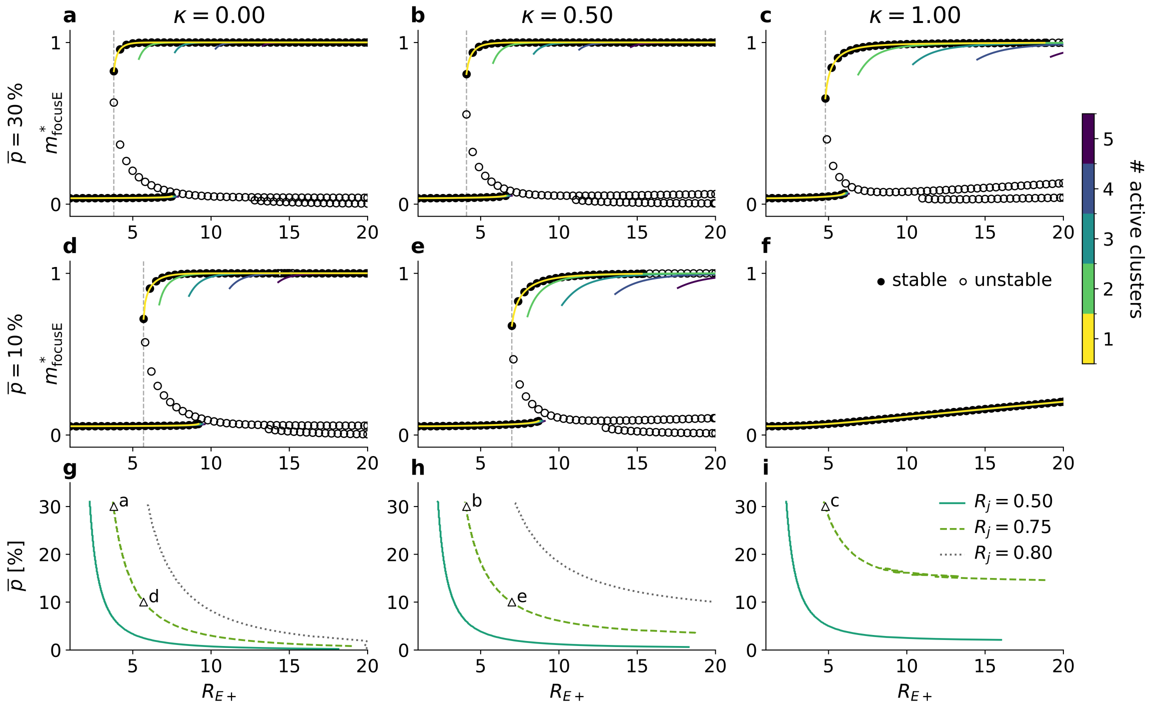


**Figure S2.** Mean-field fixed-point structure of fixed indegree construction. Compared to the Poisson multigraph case, predicted rates are closer to saturation and bifurcations appear at lower $R_{\mathrm{E+}}$. All analysis were performed with the independent closure and a root-finding solver, using the independent closure rather than the quenched closure we used in the main text. The quenched closure never converged with fixed indegree. (a–c) Predicted fixed points of the focused excitatory cluster activity $m^*_{\mathrm{focusE}}$ as a function of excitatory clustering strength $R_{\mathrm{E+}}$ at uniform connectivity $\bar{p} = 30\%$, shown for (a) $\kappa = 0$, (b) $\kappa = 0.5$, and (c) $\kappa = 1$. Colored branches correspond to solutions with different numbers of simultaneously active clusters (stable branches shown). Black markers indicate stable (filled) and unstable (open) single-active-cluster fixed points. (d–f) Same as (a–c) for $\bar{p} = 10\%$. Vertical markers indicate the first appearance of clearly separated coexisting solutions for a given $R_{\mathrm{E+}}$. (g–i) Corresponding first bifurcation boundary in the $(\bar{p}, R_{\mathrm{E+}})$ plane; metastability is possible to the right of the boundary, while only the global mode exists to the left.

### State estimation

State estimation was based on population activity traces from the binary network and comprised robust preprocessing, changepoint detection, active-set inference, and canonical post-processing. After $4 \times 10^5$ asynchronous warm-up updates, simulations were run for 900 s. The activities $m_{\alpha r}(t)$, with $\alpha \in \{\mathrm{E}, \mathrm{I}\}$ and $r = 1, \dots, N_Q$, were sampled every 12000 updates, corresponding to

$$\Delta t_{\mathrm{samp}} = 10\,\mathrm{ms}$$

and 90000 samples per simulation. State inference used only the excitatory activity vector

$$\mathbf{m}_{\mathrm{E}}(t) = \big(m_{\mathrm{E}1}(t), \dots, m_{\mathrm{E}N_Q}(t)\big),$$

because excitatory and inhibitory activities were strongly correlated. Inhibitory activities were retained for the final emission estimates.

For each run, excitatory activities were robustly scaled jointly across clusters and time,

$$\tilde{m}_{\mathrm{E}r}(t) = \frac{m_{\mathrm{E}r}(t) - \mathrm{median}_{t,r}\, m_{\mathrm{E}r}(t)}{Q_{0.95}(m_{\mathrm{E}}) - Q_{0.05}(m_{\mathrm{E}})}.$$

Joint scaling preserved relative amplitude differences between clusters. To suppress changepoints caused by flat or noisy clusters, cluster $r$ was assigned a reliability weight

$$w_r = \frac{S_r^2}{S_r^2 + \nu_r^2 + \epsilon}, \qquad S_r = \max[Q_{0.95}(m_{\mathrm{E}r}) - Q_{0.05}(m_{\mathrm{E}r}) - 2\nu_r, 0]\,,$$

where

$$\nu_r = \frac{1.4826}{\sqrt{2}}\,\mathrm{MAD}(\Delta m_{\mathrm{E}r})\,.$$

Here,

$$\mathrm{MAD}(v) = \mathrm{median}_i\,|v_i - \mathrm{median}_j\, v_j|$$

is the median absolute deviation. The factor 1.4826 makes the MAD a consistent standard-deviation estimator for Gaussian data, while $1/\sqrt{2}$ accounts for the increased variance of first differences $\Delta m_{\mathrm{E}r}(t) = m_{\mathrm{E}r}(t+1) - m_{\mathrm{E}r}(t)$.

Changepoints were detected using PELT [1], as implemented in `skchange`, with feature vector

$$\mathbf{y}_t = \big(\sqrt{w_1}\tilde{m}_{\mathrm{E}1}(t), \dots, \sqrt{w_{N_Q}}\tilde{m}_{\mathrm{E}N_Q}(t)\big)$$

and segment cost

$$C(a,b) = \sum_{t=a}^{b-1} \lVert \mathbf{y}_t - \bar{\mathbf{y}}_{a:b} \rVert_2^2\,, \qquad \bar{\mathbf{y}}_{a:b} = \frac{1}{b-a}\sum_{t=a}^{b-1} \mathbf{y}_t.$$

The PELT penalty was 10, the minimum segment length was two samples, and candidate boundaries were initially evaluated at every second samples. Boundaries were subsequently refined at full sample resolution by minimizing the local two-segment cost. Thus, the minimum segment duration was approximately 20 ms.

For each resulting interval $\mathcal{I}_\ell$, we calculated the mean scaled activity

$$\mathbf{x}_\ell = \frac{1}{|\mathcal{I}_\ell|}\sum_{t \in \mathcal{I}_\ell} \tilde{\mathbf{m}}_{\mathrm{E}}(t)$$

and inferred a binary active-set vector

$$\mathbf{z}_\ell \in \{0,1\}^{N_Q}, \qquad K_\ell = \sum_r z_{\ell r}.$$

Conditional on $K_\ell$, entries of $\mathbf{x}_\ell$ were modeled as low- or high-activity Gaussian components. An expectation–maximization procedure alternated between estimating component parameters and selecting the active set by minimizing the Gaussian negative log-likelihood plus

$$\lambda_{\text{comb}} \log \binom{N_Q}{K_\ell} + \lambda_{\text{active}} K_\ell.$$

We used

$$\lambda_{\text{comb}} = 0.1, \qquad \lambda_{\text{active}} = 0, \qquad \sigma^2_{\text{min}} = 10^{-4},$$

with at most 30 EM iterations. Candidate solutions were retained only when the active- and inactive-component means differed by at least 0.05.

Adjacent intervals with identical active sets were merged. An $A$-$B$-$A$ excursion was removed when $B$ lasted at most three samples and differed from $A$ by no more than two active-set entries, thereby suppressing weak excursions lasting approximately 30 ms or less.

Active sets were then canonicalized to remove weak spillover activity. A proposed active cluster $r$ was retained only if its episode-mean activity satisfied

$$\bar{m}_r^{(\ell)} \geq \text{median}\Big(\bar{m}_{\text{inactive}}^{(\ell)}\Big) + 3(1.4826)\ \text{MAD}\Big(\bar{m}_{\text{inactive}}^{(\ell)}\Big)\,.$$

Secondary active clusters additionally had to satisfy

$$\bar{m}_r^{(\ell)} \geq 0.5\,\bar{m}_{\text{max}}^{(\ell)},$$

and canonical active sets were restricted to at most two clusters.

Unless implied directly by the sampling interval or by the Gaussian MAD conversion, the numerical hyperparameters above were selected using pilot simulations to suppress noise-driven transitions while retaining clearly separated activity states. They were fixed before the reported analyses and applied unchanged to all networks and simulation conditions.

The final state sequence $s(t)$ assigned the canonical label to every sample in an episode. State emissions were averages of the original, unscaled activities over samples assigned to state $s$:

$$\boldsymbol{\eta}_s = \big(\langle m_{\text{E}1}\rangle_s, \ldots, \langle m_{\text{E}N_Q}\rangle_s, \langle m_{\text{I}1}\rangle_s, \ldots, \langle m_{\text{I}N_Q}\rangle_s\big)\,.$$

**Table 1. Switching statistics depend on both the clustering mixture and the distance from the multistability boundary.** All networks had $N_E = 8{,}000$, $N_I = 2{,}000$, $N_Q = 20$, and $R_{\mathrm{E}+} = 7.25$, and each simulation was analyzed over 900 s of model time. The two parameter sets correspond to the networks shown in Figs. 3 and 4. For the former, 45 independent connectivity realizations were simulated at $R_j = 0.79$; for the latter, 45 independent network realizations with three initial conditions each were simulated at $R_j = 0.75$. For the latter set, the switching rate and number of detected states were first averaged across the three initializations of each connectivity and then across connectivities. A connectivity was classified as non-switching only when none of its three initializations exhibited a state transition. "All realizations" includes networks that remained in one inferred state throughout the observation period, whereas "Switching only" excludes these non-switching networks. At fixed $R_{\mathrm{E}+}$ and $R_j$, increasing $\kappa$ was associated overall with more frequent switching, a larger repertoire of inferred states, and fewer non-switching realizations. Comparison of the two $R_j$ values further shows that a small shift in inhibitory clustering can move the networks from sparse to frequent switching. Thus, the observed metastability depends both on the clustering mixture $\kappa$ and on the location of the parameter set relative to the bifurcation.

| $\kappa$ | $N_{\mathrm{net}}$ | $N_{\mathrm{no\ switch}}$ | Switches/min | | Detected states | |
|---|---|---|---|---|---|---|
| | | | All realizations | Switching only | All realizations | Switching only |
| *Fig. 3*: $R_j = 0.79$ | | | | | | |
| 0.00 | 45 | 16 | 0.26 | 0.40 | 4.02 | 5.69 |
| 0.50 | 45 | 11 | 4.29 | 5.67 | 12.38 | 16.06 |
| 1.00 | 45 | 0 | 43.67 | 43.67 | 42.44 | 42.44 |
| *Fig. 4*: $R_j = 0.75$ | | | | | | |
| 0.00 | 45 | 19 | 0.04 | 0.07 | 1.43 | 1.74 |
| 0.10 | 45 | 11 | 0.04 | 0.05 | 1.52 | 1.69 |
| 0.20 | 45 | 7 | 0.11 | 0.12 | 1.93 | 2.10 |
| 0.30 | 45 | 2 | 0.12 | 0.12 | 2.45 | 2.52 |
| 0.40 | 45 | 2 | 0.29 | 0.30 | 3.04 | 3.13 |
| 0.50 | 45 | 2 | 0.38 | 0.40 | 3.16 | 3.26 |
| 0.60 | 45 | 1 | 0.55 | 0.56 | 3.98 | 4.05 |
| 0.70 | 45 | 1 | 0.60 | 0.61 | 3.36 | 3.42 |
| 0.80 | 45 | 3 | 0.85 | 0.91 | 3.87 | 4.07 |
| 0.90 | 45 | 2 | 0.23 | 0.24 | 2.92 | 3.01 |
| 1.00 | 45 | 2 | 1.24 | 1.29 | 5.87 | 6.09 |

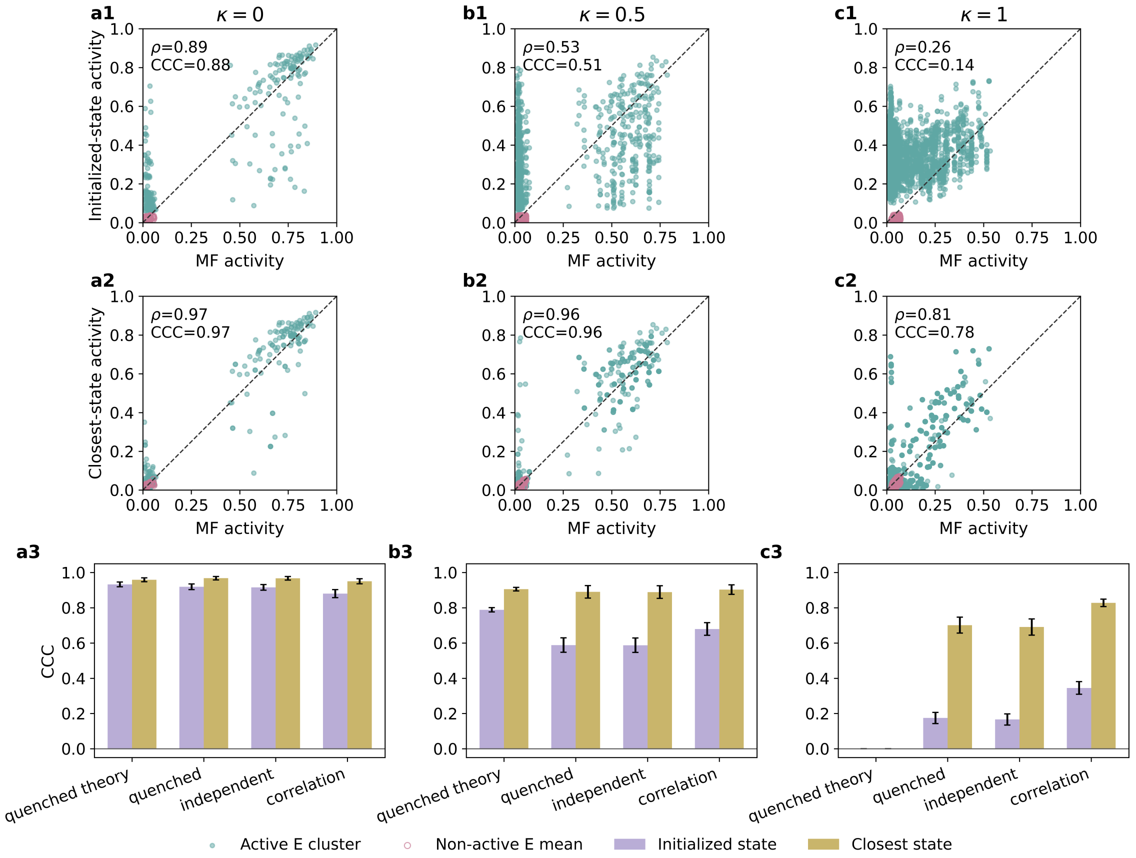


**Figure S3. Mean-field predictions for inferred state activities across clustering mixtures.** Binary network simulations ($N_E = 8{,}000$, $N_I = 2{,}000$, $N_Q = 20$, $R_{E+} = 7.25$, $R_j = 0.79$) were analyzed using the simulation ensemble and parameter set underlying Fig. 3 with 45 simulations per clustering mixture. For each inferred state, measured population activities initialized the hierarchy of mean-field solvers–independent, correlated, and quenched–using empirical connectivity moments for all three and theoretical moments for the quenched solver. Scatter plots compare activities predicted by the quenched closure using empirical connectivity moments to the observed activity of the initialized inferred state (a1–c1) or to the closest inferred state within the same network realization after convergence (a2–c2), for $\kappa = 0$, 0.5, and 1. Closest states were matched without relabeling clusters, i.e. activities remain ordered by cluster identity. Thus, the spread in the closest-state comparison reflects both the number of inferred states per network and, especially for $\kappa = 0$ where few states were detected per realization, variability across network realizations. Active excitatory clusters and the mean of inactive excitatory clusters are shown separately. Bottom panels (a3–c3) summarize prediction quality using the concordance correlation coefficient [2], $\mathrm{CCC} = \frac{2\rho\sigma_x\sigma_y}{\sigma_x^2+\sigma_y^2+(\mu_x-\mu_y)^2}$, where $x$ and $y$ denote mean-field and simulation activities, respectively, $\mu$ and $\sigma^2$ their corresponding means and variances, and $\rho$ their Pearson correlation coefficient. Bars compare quenched closure with analytic moments, quenched closure with empirical moments, independent closure, and correlated closure; error bars show the standard error across simulations.

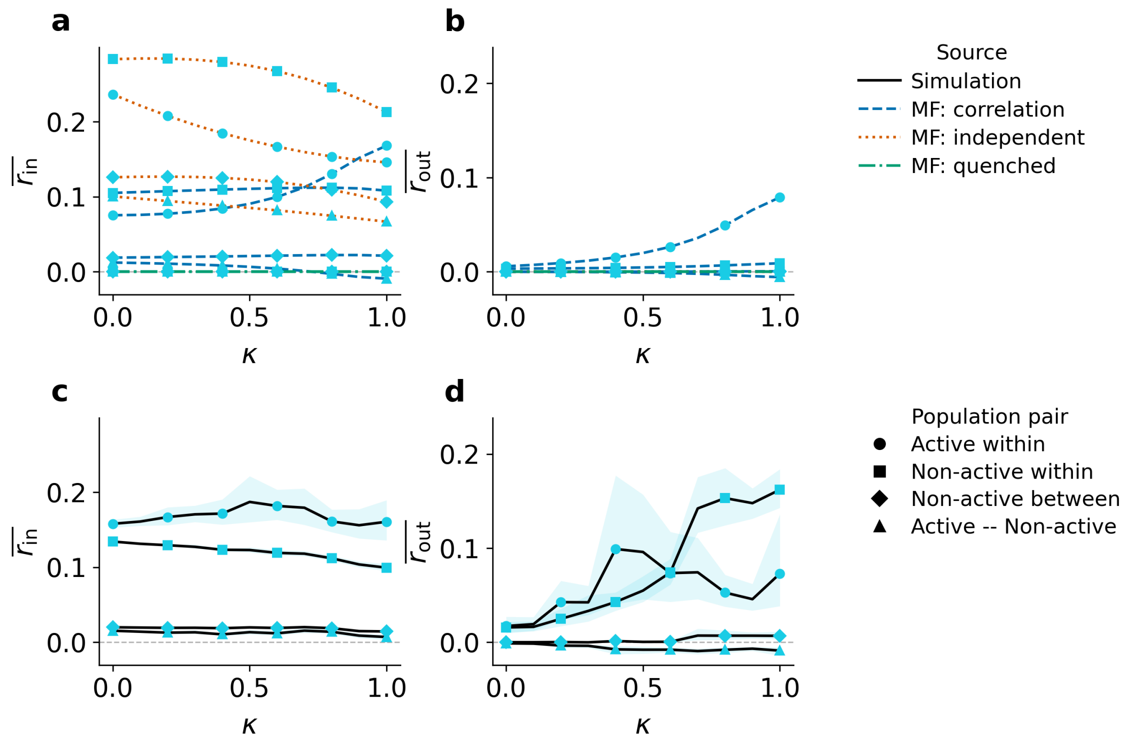


**Figure S4. Mean-field predictions for state correlations.** This figure extends the analysis of Fig. 4 using the same binary-network simulations and inferred states. Networks ($N_E = 8{,}000$, $N_I = 2{,}000$, $N_Q = 20$, $R_{\mathrm{E}+} = 7.25$, $R_j = 0.75$) were simulated for 45 independent connectivity realizations and three initial conditions per realization. After a warm-up period of $4 \times 10^5$ asynchronous updates, each run was simulated for 900 s of model time, with population activity sampled every $1.2 \times 10^4$ updates, yielding 90,000 samples per run. Statistics were computed from all inferred single-active-cluster episodes. Repeated occurrences of the same state were first averaged within each run, followed by averaging across active-cluster identities, initial conditions, and connectivity realizations. Lines show simulations and predictions from the quenched, independent, and correlated mean-field closures. Marker shape denotes the corresponding population or population-pair class. Shaded areas indicate bootstrap 95% confidence intervals across connectivity realizations. Shown as a function of $\kappa$ are average recurrent-input correlation $\bar{r}_{\text{in}}$ in theory (a) and simulation (c), and average neuronal-state correlation $\bar{r}_{\text{out}}$ in theory (b) and simulation (d).

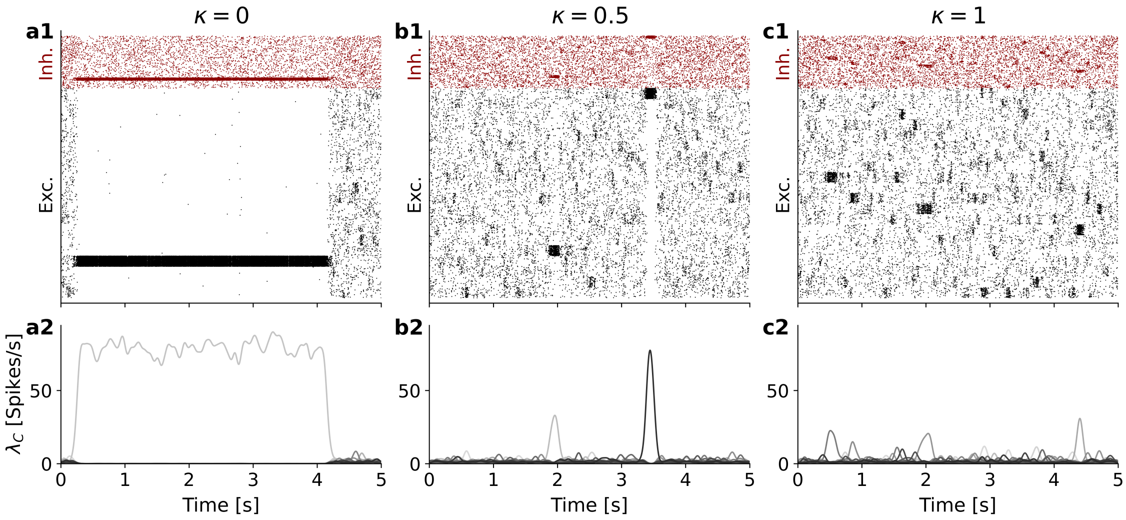


**Figure S5.** Spiking LIF networks reproduce metastable switching across clustering mixtures with fixed indegree construction ($N_E = 8000$, $N_I = 2000$, $N_Q = 20$, $R_{\mathrm{E}+} = 5.0$, $R_j = 0.78$). Top panels (**a1**–**c1**) show raster diagrams where each vertical tick represents the occurrence of an action potential. All three clustering regimes exhibit metastable switching. Bottom panels (**a2**–**c2**) show per-cluster mean firing rates $\lambda_c$, computed by Gaussian kernel convolution of spike trains ($\sigma = 50\,\mathrm{ms}$, truncated at $\pm 3\sigma$). Compared to the spiking simulations with Poisson connectivity, the rates are higher in panels (a) and (b). Compared with the Poisson-connectivity simulations, the firing rates are higher for structural and mixed clustering, indicating that the connectivity construction shifts the network operating point.
